# The proteasome and ERAD pathways can regulate hepatitis E virus pORF1 proteostasis and genome replication

**DOI:** 10.64898/2026.10.08.757567

**Authors:** Khadijah M. Abualsaoud, Frazer J.T. Buchanan, Owen Byford, Ellie M. Bartholomew, Charlotte E. Williams, Maya Kailani, Rebekah Penrice-Randal, Mark Harris, Morgan R. Herod

**Affiliations:** School of Molecular and Cellular Biology, Faculty of Health and Life Sciences and Astbury Centre for Structural Molecular Biology, University of Leeds, Leeds, LS2 9JT, UK; Centre for Proteome Research, Department of Biochemistry, Cell & Systems Biology, Institute of Systems, Molecular & Integrative Biology, Biosciences Building, Crown Street, University of Liverpool, Liverpool L69 7ZB, UK; Department of Biochemistry, Cell & Systems Biology, Institute of Systems, Molecular & Integrative Biology, Faculty of Health and Life Sciences, Biosciences Building, Crown Street, University of Liverpool, Liverpool, L69 7ZB, UK

**Keywords:** HEV, replication complex, ORF1, PCP, proteasome

## Abstract

Hepatitis E virus (HEV) is a major cause of acute viral hepatitis, with an estimated 20 million human infections annually. Its diverse host range presents a significant zoonotic threat, yet the molecular mechanisms underpinning HEV–host interactions remain poorly characterised. Here, we used transposon-based insertion screening of the HEV pORF1 polyprotein to identify a novel region within the viral PCP domain that permits HA-epitope insertion, enabling the first epitope tagging of this domain. Combined with V5 tagging of the hypervariable region (HVR), this approach enabled the generation of a replication-competent stable cell line expressing dual-tagged ORF1. Co-immunoprecipitation and TMT-based LC-MS proteomics revealed that pORF1 interacts with host proteins involved in endoplasmic reticulum-associated degradation (ERAD) and the 26S proteasome. Pharmacological inhibition demonstrated that both pathways are required for efficient HEV replication. Treatment with the FDA-approved proteasome inhibitor bortezomib altered pORF1 precursor abundance and led to loss of full-length pORF1 through enhanced proteolysis. Importantly, the kinetics of ORF1 degradation following proteasome inhibition correlated with immediate loss of HEV replication, demonstrating a functional link between pORF1 proteostasis and viral replication. Collectively, our findings identify the ERAD network as a previously unrecognised requirement for HEV replication and establish a role for the 26S proteasome in maintaining pORF1 proteostasis.

**Importance:** Hepatitis E virus (HEV) is classified in the *Hepeviridae* family, which currently comprises 10 species, and infects a wide range of animal hosts. Genetically divergent HEV strains have recently been identified within populations of rodents, bats and shrews. Thus, novel HEV zoonotic spillover events are of increasing concern. A detailed understanding of how HEV subverts host cellular machinery for replication is incomplete. Here, we demonstrate that the HEV pORF1 polyprotein interacts with components of the ERAD and proteasomal pathways. We further show that pharmacological inhibition of these networks significantly reduced HEV replication. Mechanistic analysis revealed that pORF1 homeostasis is dependent on 26S proteasomal function as bortezomib treatment resulted in aberrant HEV pORF1 proteolysis, corresponding with a reduction in genome replication. Collectively, our findings improve the understanding of HEV host-pathogen interactions and further characterise the roles of two critical pathways required during replication.

## Introduction

Hepatitis E virus (HEV) represents a major global cause of acute viral hepatitis with >20 million infections reported annually, of which ∼3.4 million are symptomatic leading to ∼ 70,000 fatalities (Li et al., 2020). HEV infection is typically acute and self-limiting (Adlhoch et al., 2016), however, in immunocompromised patients (Kamar et al., 2011) infection can become chronic and lead to cirrhosis (Dalton et al., 2009). HEV infection during pregnancy also poses a significant danger, with high incidences of maternal mortality, miscarriage, stillbirth or neonatal death (Bigna et al., 2020). Currently, there are no approved direct-acting antivirals (DAAs) to target HEV and the therapeutic agents used to date, ribavirin and pegylated interferon, have reduced efficacy due to ribavirin-resistant HEV variants (Debing et al., 2014; Nimgaonkar et al., 2018). A greater fundamental understanding of the HEV replication cycle could be key in developing future HEV DAAs or host-targeted agents (HTAs).

HEV is a member of the *Hepeviridae* family, which has recently been reclassified into two subfamilies, *Parahepevirinae* and *Orthohepevirinae*. HEV belongs to the *Paslahepevirus* genus, which comprises two species. The species *Paslahepevirus balayani* includes eight recognised genotypes (G1-G8) (Purdy et al., 2022). Among these, G1 and G2 are restricted to humans and are transmitted predominantly via the faecal-oral route, while G3 and G4 are zoonotic and can infect humans, as well as a broad range of mammals including swine, deer and wild boar (Meng, 2013; Meng, 2016).

The HEV genome is a positive-sense, single-stranded RNA of approximately 7.2 kb that encodes three open reading frames (ORFs). The genome is capped at the 5’ end with a 7-methylguanosine (m7G), and polyadenylated at the 3’ end, features that enable direct translation by host ribosomes. During replication two distinct RNA molecules are generated, the full-length genomic RNA and a shorter subgenomic RNA (sgRNA) (Fig 1A). ORF1 is translated from the genomic RNA, whereas ORF2 and ORF3 are translated from the sgRNA. ORF2 and ORF3 produce the viral capsid protein (Montpellier et al., 2018) and a small ion channel protein (Ding et al., 2017), respectively. ORF1 encodes a large non-structural polyprotein (hereafter referred to as pORF1) that is necessary and sufficient for viral RNA replication (Koonin et al., 1992). By analogy to related viruses, HEV RNA replication is thought to occur in membrane-associated replication organelles where pORF1 components are recruited together with host cell factors to make a favourable environment for viral RNA synthesis. Despite the key role of pORF1 in viral replication, the pro-viral host cell factors that are recruited by pORF1 to facilitate viral replication are relatively understudied.

**Figure 1.**
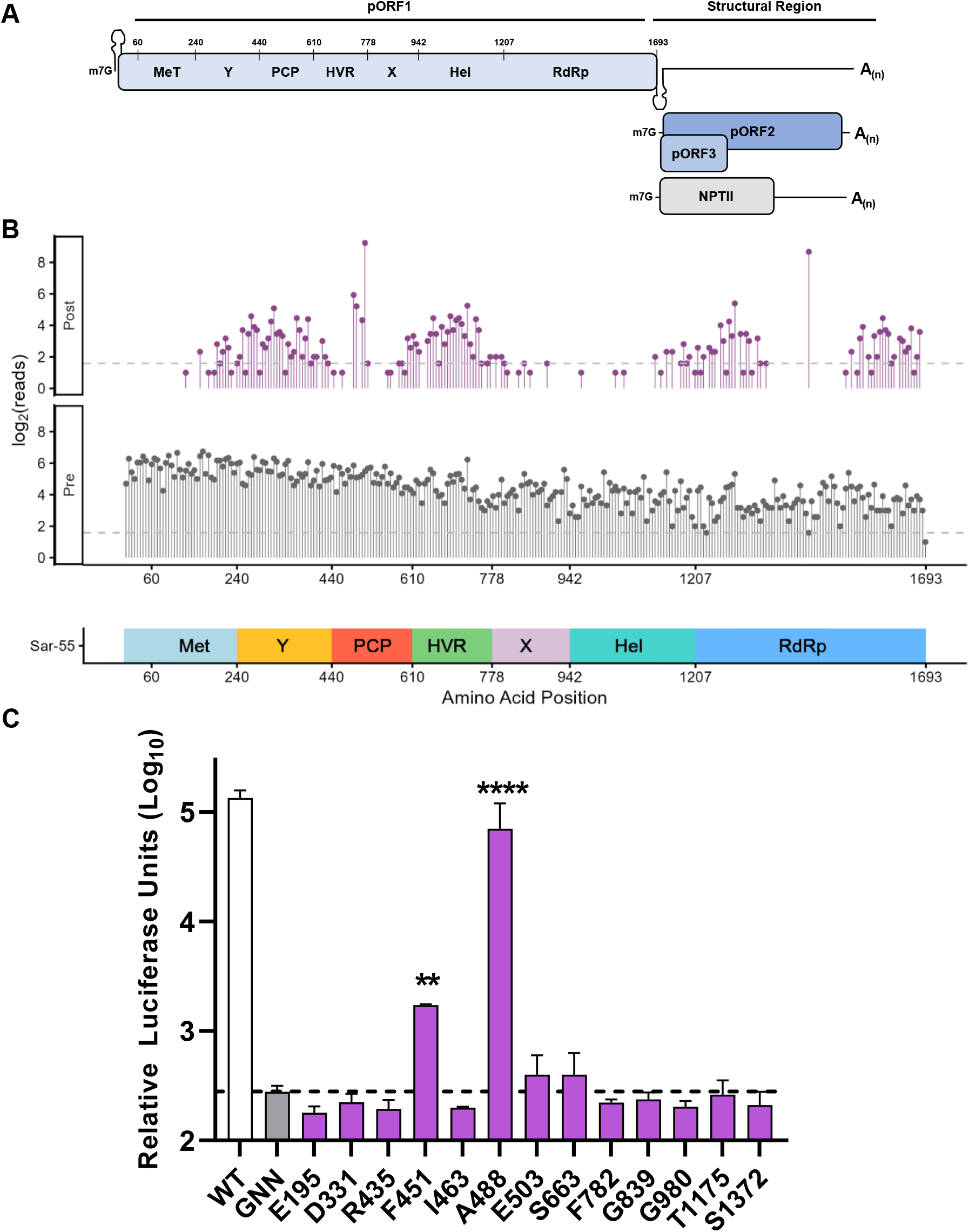
Replication of transposon insertions in Huh7 cells. (A) Schematic of the G1 HEV genome and SGR expressing NPTII in place of ORF2 and ORF3. The pORF1 polyprotein is labelled with the seven predicted functional domains and their positions within pORF1. All numbers refer to the amino acid position in the Sar55 isolate (GenBank reference AF444002). (B) Schematic of pORF1 showing log_2_ transposon insertion frequency pre- and post-selection in 10-nucleotide bins with more than 1 read. Dashed line indicates log_2_(3) reads. (C) Huh7 cells were electroporated with SGR-G1-nLuc RNA containing the individual transposon insertions, with the number indicating the amino acid residue of pORF1 after which the insertion occurred. Huh7 cells were also electroporated with wild-type (WT) and replication-defective (GNN) control SGR RNAs. Cells were harvested at 96 h post-electroporation and luciferase activity determined. Data shown represent log_10_ of mean relative luciferase activity at 96 h post-electroporation (n = 3 +/−SEM, **** P < 0.0001, ** P<0.01).

To study pORF1 host-cell interactions, many investigations have focused on expressing individual functional domains separately, such as the methyltransferase (Met), helicase (Hel), hypervariable region (HVR), and RNA-dependent RNA polymerase (RdRp) (Subramani et al., 2018; Kanade et al., 2019). In contrast, only one study to date has incorporated a tag into the full-length pORF1 within a subgenomic replicon (SGR) to explore host factors involved in HEV replication (Ju et al., 2023). Whilst these findings increased the understanding of HEV restriction factors, the study identified few pro-viral host proteins, possibly because of the SGR genetic complementation system used. The current knowledge gap surrounding pORF1 function and host factor interactions represents a major obstacle to understanding HEV replication and developing targeted antivirals.

Here, we have identified pro-viral host factors that interact with HEV pORF1. By employing transposon mutagenesis, we developed an epitope-tagged permissive HEV SGR to generate a chronic model of HEV replication. Using pORF1 co-immunoprecipitation coupled with tandem mass tag (TMT) based mass spectrometry we identified pro-viral host factors that support viral replication. This revealed enrichment of proteins associated with the proteasome and the endoplasmic reticulum-associated protein degradation (ERAD) pathway that transports ubiquitinated proteins to the proteasome for processing. Using specific proteasome inhibitors, we confirmed that HEV replication is dependent upon a functional proteasomal pathway. Importantly, the observed reduction in viral replication corresponded with a loss of native pORF1 products. Similarly, with the use of HA-15 (HSPA5 inhibitor), CB-5083 (p97 inhibitor) and TAK-243 (ubiquitin inhibitor), which target differing stages of the ERAD pathway, we demonstrate a requirement for a functioning ERAD network during HEV replication. Together these findings suggest that HEV depends on a functional ERAD pathway to direct pORF1 to the proteasome and maintain long-term pORF1 stability during replication.

## Results

### Transposon mutagenesis of the HEV ORF1 polyprotein

Multiple studies have used random transposon-mediated mutagenesis to probe the genomes of positive-strand RNA viruses for sites that can tolerate the insertion of small exogenous sequences (Teterina et al., 2011; Herod et al., 2015; Perry et al., 2018). Previous studies with HEV have successfully used this approach with a G3 isolate; however, this study identified only sites that could tolerate epitope insertions in the HVR region which retain replicative capacity (Szkolnicka et al., 2019; Buchanan et al., 2023). To expand on this genetic toolkit, we employed a G1 SGR for random transposon mutagenesis of ORF1.

For this a plasmid containing the HEV genotype 1 SGR expressing nano-luciferase (SGR-G1-nLuc, Sar55 isolate) was used to generate an SGR library containing insertions solely in the pORF1 polyprotein coding sequence. To facilitate generation of the library the SGR plasmid required modification to allow restriction enzyme digestion for sub-cloning. Site-directed mutagenesis was therefore used to introduce a unique MfeI restriction site at the end of pORF1 and a NotI site within pORF1, was removed via introduction of a silent nucleotide change. Neither of these changes significantly affected SGR replication (Figure S1A).

Subsequently, the modified ORF1 region was sub-cloned into pUC19 for transposon mutagenesis, to ultimately generate the transposon-modified SGR plasmid library. This resulted in a library of 15-nucleotide (nt) insertions spanning HEV pORF1. Each 15 -nt insertion consisted of 10 -nt derived from the transposon insertion (including a unique NotI restriction enzyme site) and 5-nt derived from the target DNA directly upstream of the insertion site, which is duplicated during the transposition event. To permit selection, the transposon modified SGR plasmid library was introduced into an SGR expressing neomycin phosphotransferase II (NPTII) in place of nano-luciferase (termed SGR-G1-NPTII) (Figure 1A).

To identify replication permissive insertion sites, *in vitro* transcribed RNA from the transposon-mutated SGR library was electroporated into Huh7 cells in duplicate alongside the wild-type (WT) SGR-G1-NPTII and replication-defective control SGR containing a point mutation within the viral polymerase (GNN), which acted as a control for input translation. Electroporated cells were either seeded into 6-well plates for colony formation assay or seeded into dishes and maintained as a polyclonal cell line for RNA isolation and sequencing. At 24 h post-electroporation, all cells were selected with G418 and maintained under selection for 14 days.

In the colony formation assay, the WT SGR yielded an average of 290 colonies per 1 μg of RNA (Figure S1B). As anticipated, the GNN control SGR RNA produced no colonies. In comparison cells electroporated with the transposon-modified library SGR RNA yielded on average 42 (+/−2) colonies per 1 μg of RNA. This was expected, as not all insertions within the genome are compatible with replication.

Subsequently, RNA was extracted from the Huh7 cells maintained in plates as a polyclonal cell line. Three overlapping amplicons spanning the pORF1 region were generated by RT-PCR. To provide a holistic overview of transposition within pORF1 paired-end Illumina sequencing was employed on both the pre-selected plasmid DNA library and post-selected SGR RNA sequences. Bioinformatic analysis was conducted in line with previous viral transposon mutagenesis pipelines (Heaton et al., 2013; Fulton et al., 2018).

The pre-selected library had a broad distribution of potential transposition sites (Figure 1B). Following selection of replication-competent genomes, fewer transposition sites were identified across the genome (Figure 1B), with clusters of transpositions in the Y-domain, HVR and sections of the RdRp. From the post-selected insertion sites, 13 were selected for downstream analysis based on frequency of insertion and even distribution across pORF1. To achieve this we employed a previously described HEV SGR-G1-nLuc in place of the viral structural proteins (Buchanan et al., 2023). Monitoring luciferase production allows indirect quantification of viral replication. This SGR was modified to individually introduce selected transposon insertions. RNA transcripts were generated and electroporated into Huh7 cells alongside the WT SGR and replication-defective GNN control (Figure 1C).

As expected, the WT SGR-G1-nLuc demonstrated a >100-fold increase in luciferase activity over the duration of the experiment compared to the replication-defective control (Figure 1C). Several individual transposon clones showed luciferase activity above the replication-defective control, although this increase was significant only for insertion sites after amino acids F451 and A488. The insertion site after amino acid position A488 within the PCP domain demonstrated the highest luciferase activity, with a non-significant ∼2-fold reduction in luciferase activity compared to the WT SGR. This was one of the most abundant insertion sites in the post-selected library. Thus, A488 within the G1 pORF PCP domain was identified as a site that tolerates small genetic insertions while maintaining WT replicative capacity.

### Generation of epitope-tagged HEV constructs

Identification of replication-competent insertion sites suggests potential locations for insertion of alternative exogenous sequences such as epitope tags. One frequently used small epitope tag, haemagglutinin (HA: amino acid sequence YPYDVPDYA), was chosen to replace the P487 and A488 residues. This strategy was chosen for two reasons. Firstly, although it removed both P487 and A488, the terminal alanine residue of the HA tag recapitulated A488, allowing a smaller insert that was likely to increase the chance of replication competency. Secondly, this insertion site is within the PCP domain and would allow investigation of a key viral domain which has no agreed biological function. Replication of this new epitope-tagged SGR (termed PCP^HA^) was assessed in Huh7 cells along with the relevant controls as before (Figure 2A and S2A). The WT control demonstrated a >100-fold increase in luciferase activity across the time course compared to the replication-defective GNN control. In comparison, the SGR-G1-PCP^HA^ demonstrated a non-significant ∼2.5-fold lower luciferase activity at 96 h.

**Figure 2.**
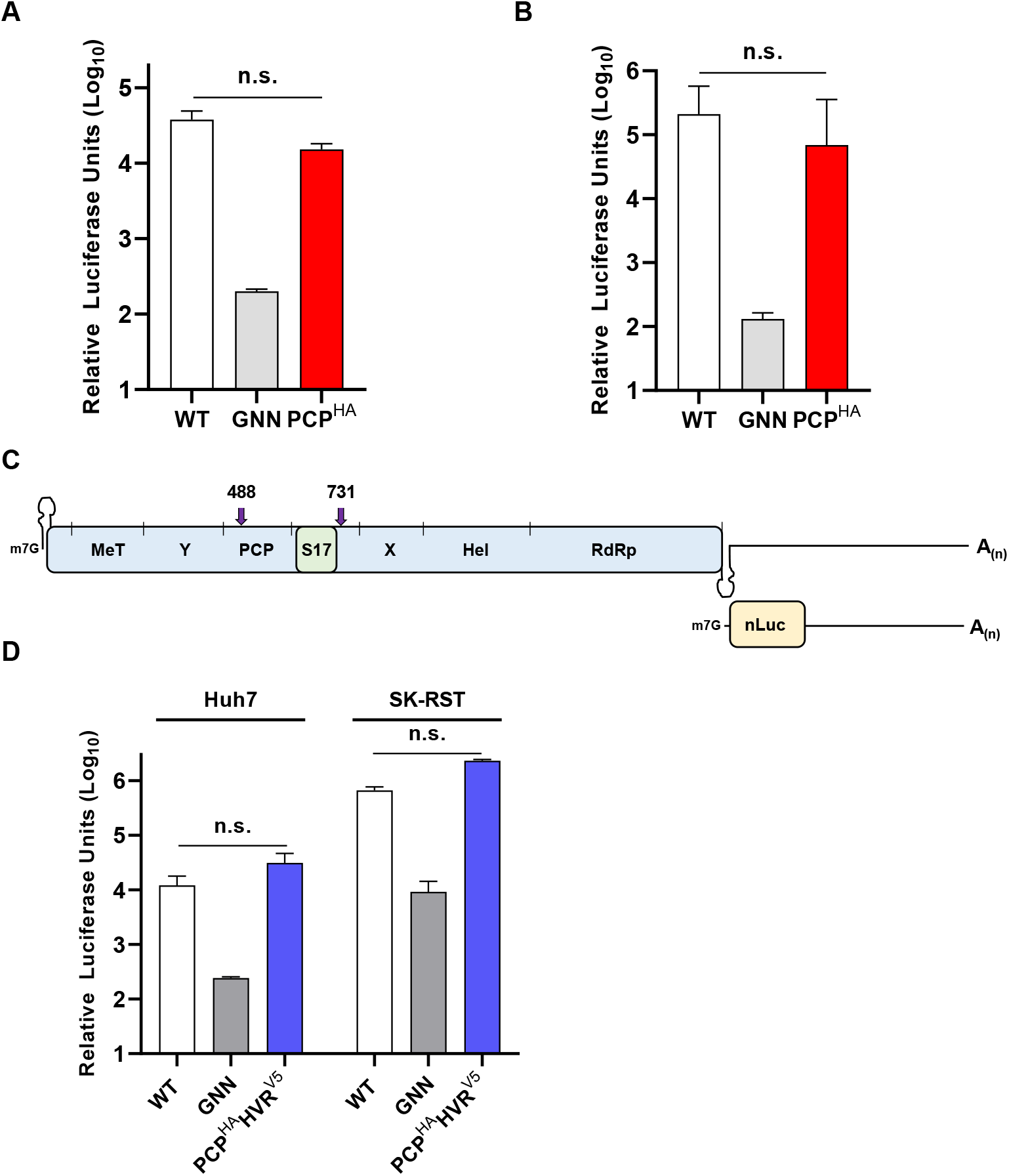
Replication of epitope-tagged HEV SGRs. Huh7 cells were electroporated with (A) SGR-G1-PCP^HA^ or (B) SGR-G3-PCP^HA^ alongside wild-type (WT) and replication-defective (GNN) controls. Cells were harvested at 24 h intervals across a 120 h time course, and luciferase activity was determined. Data shown as log_10_ of mean relative luciferase units at 96 h post-electroporation (n = 3 ± SEM). (C) Schematic of the G3 HEV SGR containing S17 insertion and PCP^HA^ tag and HVR^V5^ tag. (D) Huh7 and SK-RST cells were electroporated with the dual-tagged SGR-G3-PCP^HA^HVR^V5^ alongside wild-type (WT) and replication-defective (GNN) controls. Cells were harvested at 24 h intervals across a 120 h timecourse and luciferase activity determined. Data are shown as log_10_ of mean relative luciferase units at 96 h post-electroporation (n = 3 ± SEM).

To investigate whether the PCP^HA^ insertion was compatible with replication across genotypes we employed a G3 SGR. We inserted the PCP^HA^ epitope in the previously described 83-2/P6^80^ SGR (Buchanan et al., 2023), selected due to its high replicative capacity, to generate SGR-G3-PCP^HA^ and tested replication alongside relevant controls as before (Figure 2B and S2B). As with the SGR-G1-nLuc the SGR-G3-nLuc control demonstrated a >100-fold increase in luciferase activity across the time course. Similarly, there was no significant difference in luciferase activity between the control and tagged SGR at 96 h. These data suggest that HA-epitope insertion at this position in PCP does not significantly affect viral replication across multiple genotypes.

Following characterisation of the individual PCP^HA^ insertion, we generated a dual-tagged G3 SGR (Figure 2C) to test replication competency in combination with the previously published HVR epitope insertion after amino acid 731 (Pierce et al., 2023).

Replication of this new dual epitope-tagged SGR (termed SGR-G3-PCP^HA^HVR^V5^) was assessed along with the relevant controls in both Huh7 (human) and SK-RST (porcine) cell lines (Figure 2D and S2C/D). The WT SGR showed a >100-fold increase in luciferase activity across the time course in both cell lines compared with the replication-defective GNN control. Compared to the WT control, the SGR-G3-PCP^HA^HVR^V5^ demonstrated no significant difference in luciferase activity at 96 h. Overall, these data suggest that incorporation of both the HA and V5 tags within pORF1 did not impair replication of a G3 SGR (Figure 2D and S2C/D).

### Generation of a stable cell line harbouring an epitope dual-tagged G3 SGR

To identify viral-host protein interactions involved in HEV RNA replication, we generated a modified SGR-G3-PCP^HA^HVR^V5^ in which the nLuc reporter was exchanged for NPTII. In vitro transcribed RNA from this SGR was electroporated into Huh7 cells and used to establish a stable cell line harbouring the SGR by selection in the presence of G418 (termed Huh7-SGR-G3-HA/V5). The cell line was validated by detection of NPTII (expressed from the viral sub-genomic RNA) in cell lysate by western blot (Figure S3).

To determine the subcellular localisation of HEV tagged proteins in this cell line, we performed immunofluorescence analysis of PCP^HA^ and HVR^V5^. Confocal microscopy revealed that PCP^HA^ was predominantly distributed as cytoplasmic puncta with enrichment in the perinuclear region. The HVR^V5^ exhibited a similar punctate pattern (Figure 3A). Co-localisation between PCP^HA^ and HVR^V5^ was evident, which was most apparent in discrete foci surrounding the perinuclear subcellular region. This was consistent with line-scan analysis, which showed that the PCP^HA^ and HVR^V5^ peaks were in a close coincidence (Figure 3B). Quantification, utilising Mander’s overlap co-efficient, of 10 individual cells, revealed a ∼70% co-localisation of PCP^HA^ with HVR^V5^, whereas up to ∼60% of HVR^V5^ co-localised with PCP^HA^ (Figure 3C). These findings indicate that the HEV pORF1 protein is expressed within subcellular compartments, suggesting co-localisation between the PCP and HVR pORF1 domains during viral replication.

**Figure 3.**
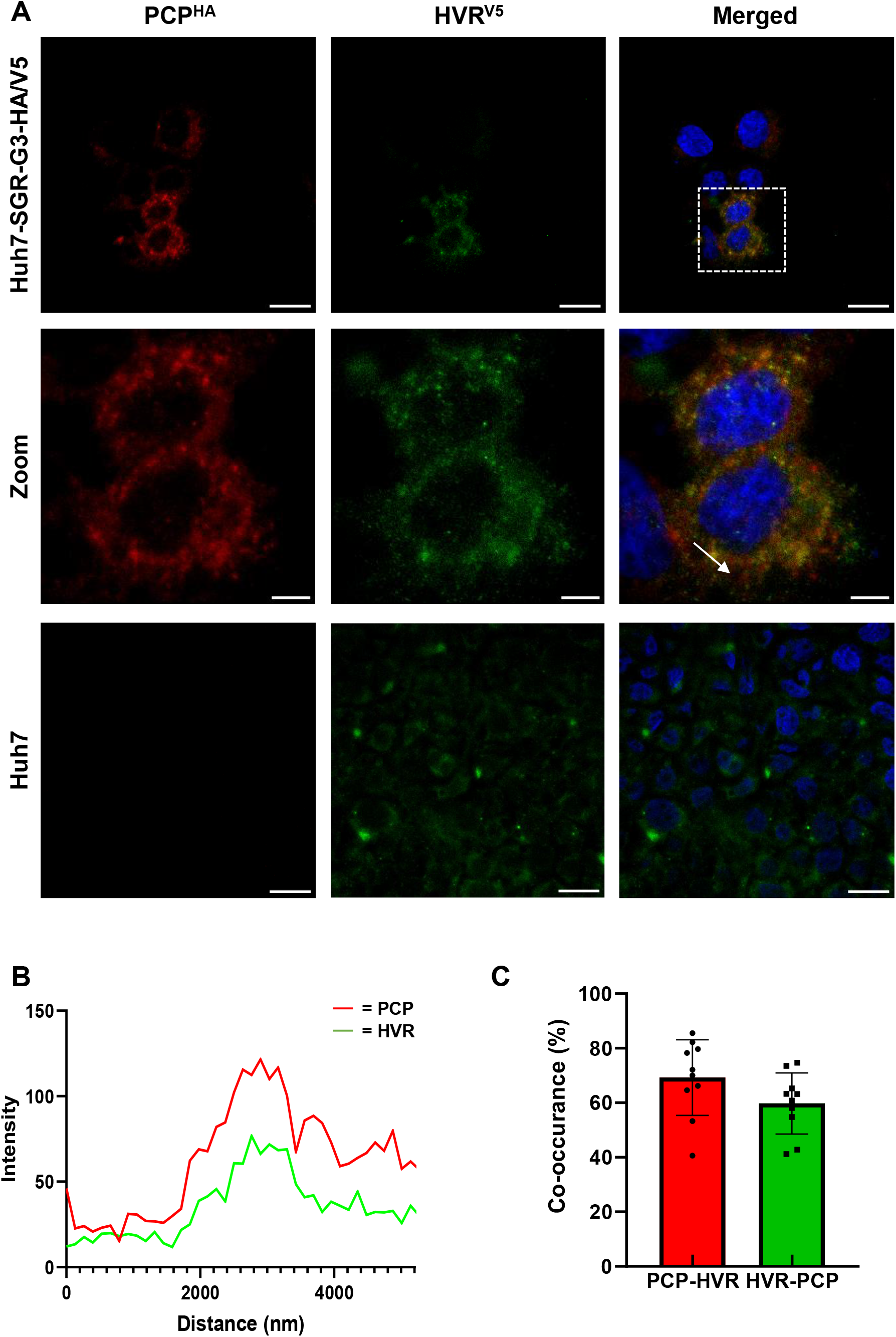
Immunofluorescence analysis of stable cell line expressing epitope-tagged G3 SGR. (A) Naïve Huh7 cells or Huh7-SGR-G3-HA/V5 were fixed and stained with anti-HA to detect PCP (red) and anti-V5 to detect HVR (green). Nuclei were counterstained with NucBlue and images were captured on a Zeiss LSM880 microscope. A zoomed-in region with overlapping puncta is shown in the merged image (arrow). Scale bar shown on the representative Huh7 and Huh7-SGR-G3-HA/V5 images are 20 µm, whereas zoomed images scale bar shown are 5 µm. (B) Line-scan for an area of interest (indicated by the white arrow) demonstrates similar intensity peaks for both anti-HA and anti-V5. (C) Quantitative co-localisation analysis was performed using Mander’s overlap co-efficient analysis and expressed as a percentage of PCP-HVR (red) and also HVR-PCP (green) (n = 10 +/-SEM).

Next, immunofluorescence analysis was performed to examine the localisation of pORF1 and dsRNA (a marker for active RNA replication of positive-sense RNA viruses). Interestingly, it appeared that the majority of dsRNA puncta observed within the Huh7-SGR-G3-HA/V5 cells did not colocalise with pORF1 as detected with HVR^V5^ (Figure 4A). Nevertheless, specific dsRNA puncta were observed to have co-localisation with HVR^V5^. Line scan analysis was performed to highlight differing dsRNA puncta whereby HVR^V5^ was either present or absent (Figure 4B). Consistently, quantification utilising Mander’s overlap co-efficient, of 10 individual cells, revealed a ∼20% co-localisation of dsRNA puncta with HVR^V5^, whereas up to ∼60% of HVR^V5^ puncta co-localised with dsRNA (Figure 4C). Collectively, these data suggest that the majority of HEV pORF1 is localised to subcellular regions that contain dsRNA.

**Figure 4.**
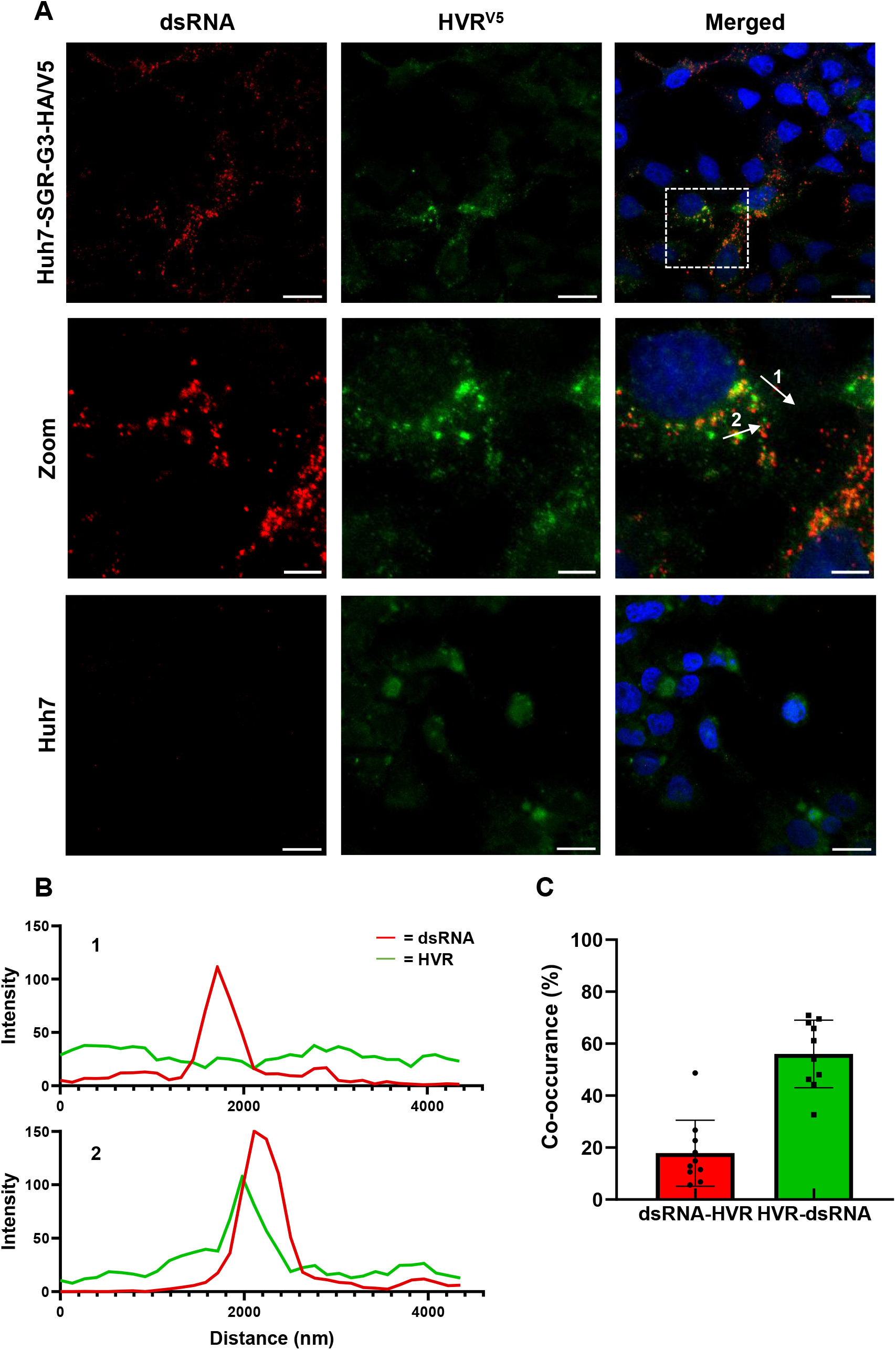
Immunofluorescence analysis of dsRNA in the Huh7-SGR-G3-HA/V5 stable cell line. (A) Naïve Huh7 cells and Huh7-SGR-G3-HA/V5 were fixed and stained with anti-dsRNA (red) and anti-V5 (green) to detect the HVR region of ORF1. Nuclei were counterstained with NucBlue. Scale bar shown on the representative Huh7 and Huh7-SGR-G3-HA/V5 images are 20 µm, whereas zoomed images scale bar shown are 5 µm. (B) Line-scan for an area of interest demonstrates intensity peaks for both anti-dsRNA and anti-V5. (C) Quantitative co-localisation analysis was performed using Mander’s overlap co-efficient analysis and expressed as a percentage of dsRNA-HVR (red) and also HVR-dsRNA (green) (n = 10 +/-SEM).

### Co-immunoprecipitation and quantitative proteomic analysis of pORF1 proteins from Huh7-SGR-G3-HA/V5 cell line

We were unable to detect pORF1 by western blotting of Huh7-SGR-G3-HA/V5 lysates with either anti-HA or anti-V5 antibodies, presumably due to the low levels of HEV translation in these cells. However, the Huh7-SGR-G3-HA/V5 stable cell line permitted a proteomic analysis of the host proteins that interact with pORF1 components.

Lysates from Huh7-SGR-G3-HA/V5 and naive Huh7 control cells were subjected to either anti-HA or anti-V5 immunoprecipitation (IP) using anti-HA or anti-V5 nanobodies conjugated to magnetic beads. Successful IP was first confirmed by western blotting the samples with both anti-HA and anti-V5 antibodies (Figure 5A and 5B). HA and V5 reactive products of apparent molecular weights (MWs) ∼200 kDa and ∼100 kDa were detected in both anti-V5 and anti-HA IP. These MWs are consistent with the predicted masses of full-length (uncleaved) pORF1 and a C-terminally truncated or proteolytically cleaved pORF1 product.

**Figure 5.**
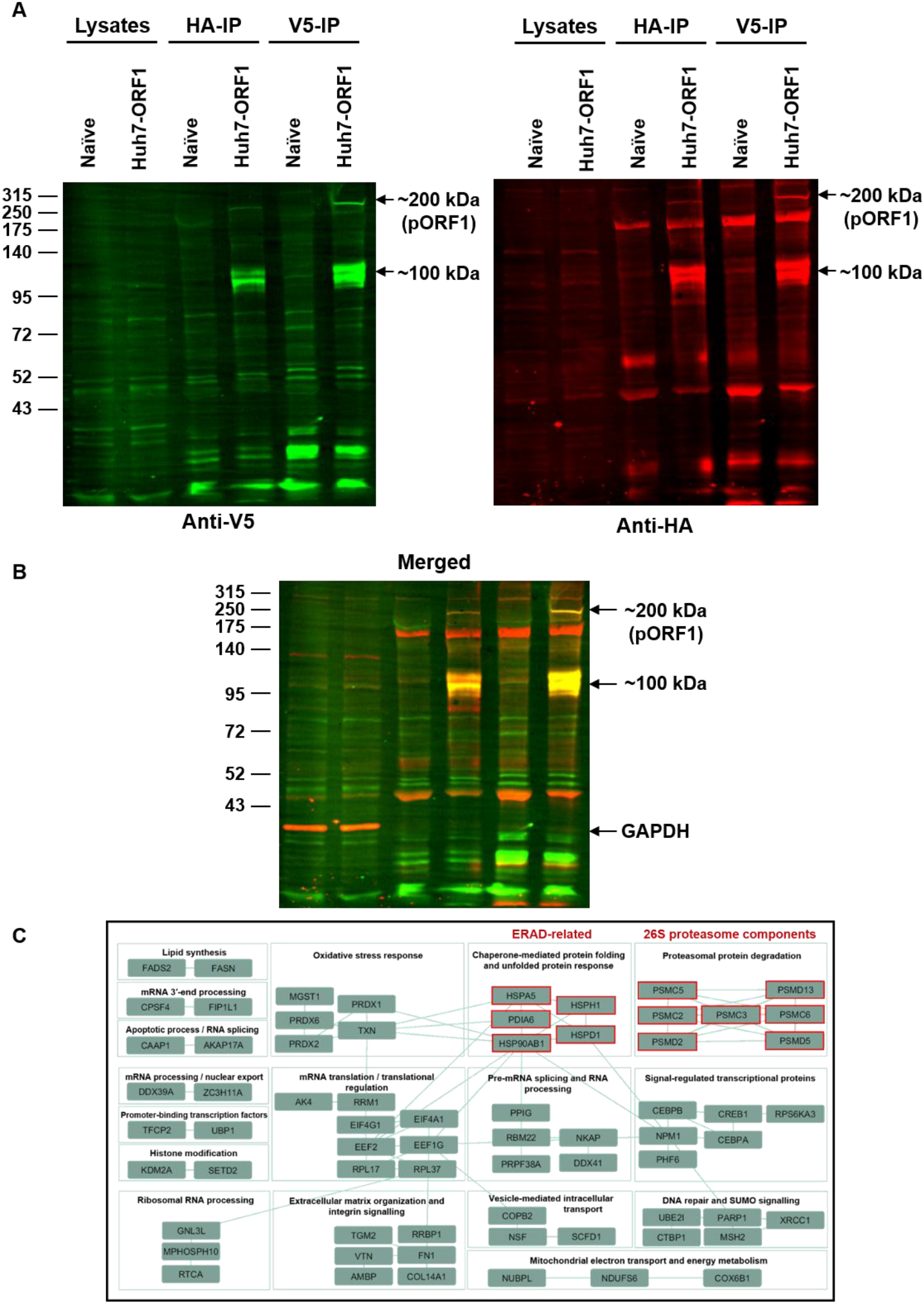
Co-immunoprecipitation and quantitative proteomic analysis of pORF1 proteins from Huh7-SGR-G3-HA/V5 cell line. (A) Lysates from naïve Huh7 control or Huh7-SGR-G3-HA/V5 cells were subjected to IP using anti-HA or anti-V5 nanobodies. Cell lysates and IP samples were analysed by western blotting with anti-V5 or anti-HA. The full-length pORF1 (∼200 kDa) and two shorter pORF1-related proteins (∼100 kDa) are indicated. Molecular weight markers (kDa) are indicated on the left of the anti-V5 blot. (B) Merged image of both anti-V5 and anti-HA blots. Regions revealing yellow are overlapping signals from both blots. GAPDH as a loading control. (C) Proteins meeting the significance criteria (false discovery rate FDR-pass, p-value <0.05 and log2FC >1) were selected for subsequent gene ontology enrichment analysis. GO enrichment analysis based on Cytoscape software with ClueGO was used to connect proteins linked via cellular pathways and GO molecular function.

The anti-HA IP sample was subsequently subjected to tandem mass tag (TMT) based liquid chromatography-mass spectrometry (LC-MS) analysis to identify cellular proteins enriched within the Huh7-SGR-G3-HA/V5 samples. From three biological repeats of the anti-HA IP, a total of 128 proteins exhibited a statistically significant enrichment (defined as having a log fold-change of > 1 alongside a P value < 0.05) in the Huh7-SGR-G3-HA/V5 compared to the naive Huh7 control cells (shown in a volcano plot: Figure S4). Within the LC-MS analysis, 10 individual unique HEV ORF1 peptides were detected from the Huh7-SGR-G3-HA/V5 stable cell line distributed across the polyprotein.

Multiple proteins identified in the LC-MS analysis were key components of the 19S regulatory component of the 26S proteasome, a central complex in the ubiquitin-proteasome system. These include the non-ATPase subunits PSMD13, PSMD5 and PSMD2, and the ATPase-containing base subunits PSMC2, PSMC3, PSMC5, and PSMC6, indicating direct engagement with the proteasomal machinery (Figure 5C).

Intriguingly, the LC/MS analysis also identified multiple members of the ERAD pathway, a cellular process that translocates proteins to the 26S proteasome for degradation (Vembar and Brodsky, 2008). In particular, HSPA5 (also named GRP78/BiP) that acts as a master regulator for identifying and processing of misfolded proteins (Rehati et al., 2023; Liu et al., 2023), HSPH1 and HSPD1 which are critical co-chaperones that recognise misfolded proteins for translocation to the proteasome (Saibil, 2013; Tian et al., 2021; Zhang et al., 2024), HSP90AB1 which stabilises misfolded proteins and facilitates proteasomal degradation (Haase and Fitze, 2016; Wang et al., 2026), UCHL1 which is important for recycling of ubiquitin to ensure a surplus exists for targeting of misfolded proteins (Mi and Graham, 2023) and PDIA6 which is a reductase in the ER lumen required to break disulphide bonds within misfolded substrates (Eletto et al., 2014). The 128 significantly enriched proteins were subject to gene ontology (GO) analysis to identify the biological relevance of the pORF1 cellular interactors. GO enrichment analysis was performed using Cytoscape with the ClueGO plugin (Figure 5C). As expected, multiple host proteins from the proteasomal protein degradation pathway were highlighted as pORF1 interactors; however, other cellular processes were revealed, included pre-mRNA splicing and RNA processing, mRNA translation and oxidative stress (Figure 5C).

### Pharmacological inhibition of the proteasome or ERAD pathway restricts HEV replication

The LC-MS analysis suggested a strong interaction between HEV pORF1 and 26S host proteasomal components (Figure 5). These observations agree with a recent study suggesting that proteasomal processing of pORF1 generates a viral-derived peptide that stimulates liver fibrosis (Zhang et al., 2025). Furthermore, pharmacological inhibition of the proteasome with MG132 has been shown to prevent HEV replication (Karpe and Meng, 2012). However, MG132 has also been demonstrated to have off-target effects and inhibit other proteases such as calpains and cathepsins (Kisselev and Goldberg, 2001; Xu et al., 2015). To confirm these observations, we first determined the sensitivity of viral genome replication to pharmacological inhibition of the proteasome using the more selective and reversible inhibitor bortezomib (Boccadoro et al., 2005). Huh7 cells were electroporated with SGR-G3-nLuc, and then subsequently treated with increasing concentrations of bortezomib (Figure 6A) (Boccadoro et al., 2005; Chen et al., 2011). Simultaneously, cell viability was measured by MTS assay to separate cytotoxicity from inhibition of genome replication (Figure 6A). Bortezomib treatment resulted in a dose-dependent reduction in HEV genome replication compared to the DMSO (vehicle-only) control, with the highest concentration tested significantly reducing replication by ∼80% (Figure 6A). All bortezomib concentrations tested had minimal impact on cellular viability (red dotted line; ∼20% reduction). To examine whether the reliance is conserved across differing genotypes, we examined the effect of bortezomib on SGR-G1-nLuc (Figure 6A). Similarly, a reduction in luciferase activity of approximately 80% was observed for cells treated with 1000 nM bortezomib. These data confirm previous observations suggesting that the proteasome is a key component required for HEV genome replication and that this reliance is conserved across different genotypes.

**Figure 6.**
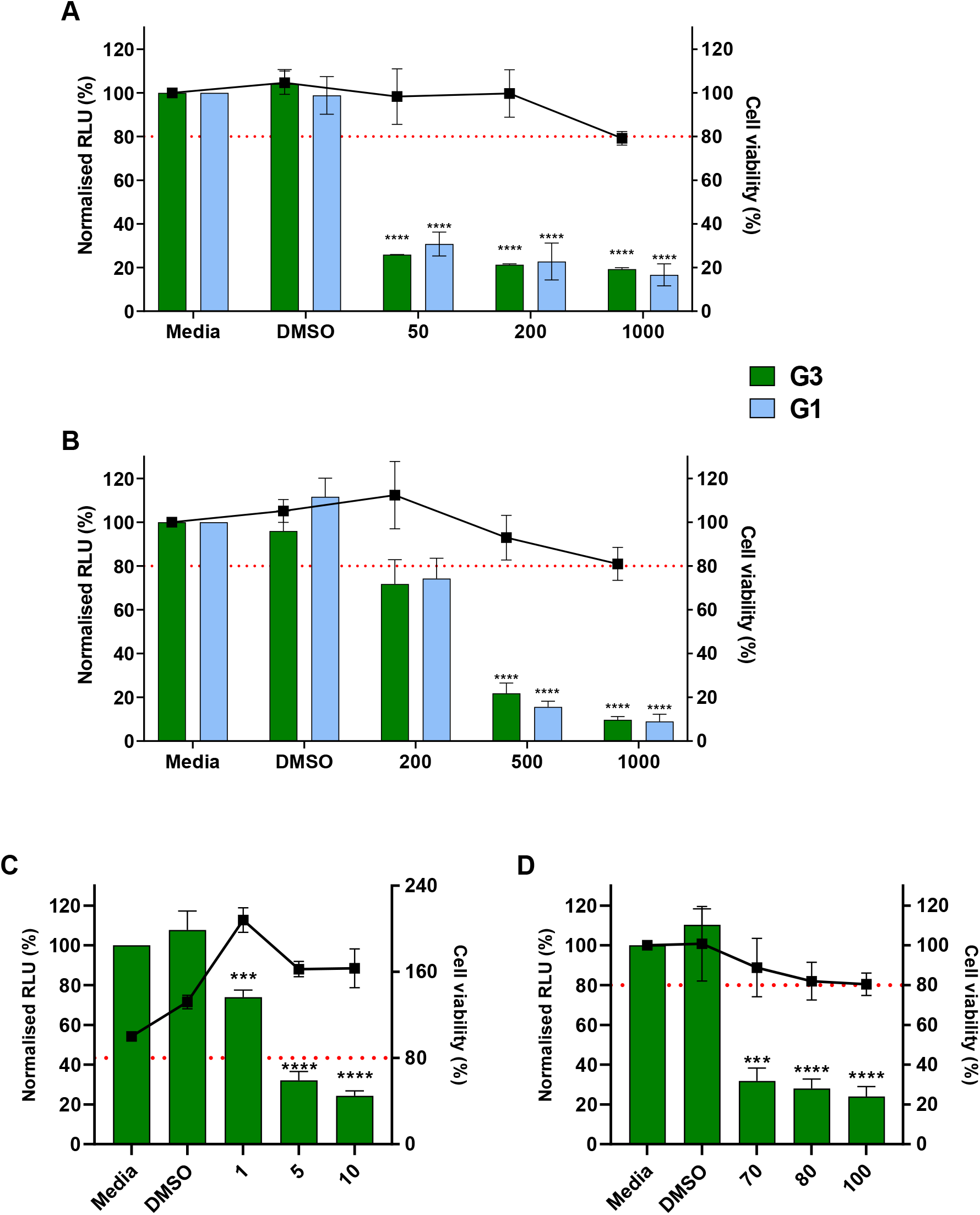
Treatment with the proteasomal inhibitor bortezomib and ERAD inhibitors reduces HEV replication. (A) Huh7 cells were electroporated with SGR-G3-nLuc or SGR-G1-nLuc RNA before the addition of bortezomib at 24 h post-electroporation. Medium-only and DMSO vehicle controls were included, and cell viability was assessed by MTS analysis. Supernatants were collected 48 h post-electroporation and luciferase activity was measured. Data shown represent a normalisation of mean relative luciferase activity (green bars, left y-axis) and MTS viability (black line; right y-axis) compared to the medium-only control. The red dotted line represents 80% viability as a non-cytotoxic cut-off (right y-axis). (B) Huh7 cells were electroporated with SGR-G3-nLuc or SGR-G1-nLuc RNA before the addition of CB-5083. Huh7 cells were electroporated with SGR-G3-nLuc RNA before the addition of HA-15 (C) and TAK-243 (D) at 24 h post-electroporation. Media-only and DMSO vehicle controls were included. Supernatants were collected 48 or 72 h post-electroporation and luciferase activity was determined. Data shown represent a normalisation of mean relative luciferase activity to media control at the respective time post-electroporation (n = 3 +/-SEM, **** P < 0.0001, *** P<0.001, ** P<0.01, * P<0.05, n.s. non-significant).

It has previously been proposed that the C-terminal region of HEV pORF1 is ubiquitinated, and this is required for proteasomal processing of pORF1 (Zhang et al., 2025). Protein within the endoplasmic reticulum (ER) that are destined for proteasomal degradation can undergo dislocation into the cytoplasm and ubiquitination by the ERAD pathway. To investigate the potential role of the ERAD pathway within HEV replication, we utilised a range of specific pharmacological inhibitors of the pathway. CB-5083, a potent and specific inhibitor of the p97 ATPase (also named VCP), which prevents the translocation of misfolded proteins from the ER (Figure 6B) (Le Moigne et al., 2017), HA-15, a specific inhibitor which binds HSPA5 to prevent its ATPase activity (Wu et al., 2020; Liu et al., 2023) (Figure 6C) and TAK-243, which is known to target the E1 ubiquitin-activating enzyme (UBA1), preventing the early stages of the ubiquitin cascade (Arakawa et al., 2024) (Figure 6D). Huh7 cells were electroporated with SGR-G3-nLuc and subsequently treated with increasing concentrations of these three inhibitors. Simultaneously, cell viability was measured by MTS assay to separate cytotoxicity from inhibition of genome replication (Figure 6B-D).

Treatment with all three inhibitors reduced HEV genome replication in a dose-dependent manner, and these concentrations that had minimal effects on cell viability. To confirm that this pathway was important across multiple HEV genotypes, the inhibitor CB-5083 was also tested against the SGR-G1-nLuc (Figure 6B). Consistent with the SGR-G3-nLuc, there was a dose-dependent reduction in replication of the G1 SGR upon treatment with CB-5083. Taken together, these data suggest that the ERAD pathway, in association with the proteasomal network is required for HEV replication, and that this requirement is conserved between genotypes 1 and 3.

### The proteasome is required for functional pORF1 proteolysis

Proteasomal inhibition has previously been shown to prevent proteolysis of pORF1 (Zhang et al., 2025). However, these studies did no clarify how HEV pORF1 is directed to the host proteasome for proteolysis while preventing complete pORF1 degradation. To confirm the impact of proteasomal inhibition on pORF1 proteolysis, Huh7-SGR-G3-HA/V5 cells were treated with 1000 nM bortezomib, followed by anti-HA or anti-V5 IP for PCP^HA^ or HVR^V5^ and western blotted with anti-HA or anti-V5 antibodies (Figure 7).

**Figure 7.**
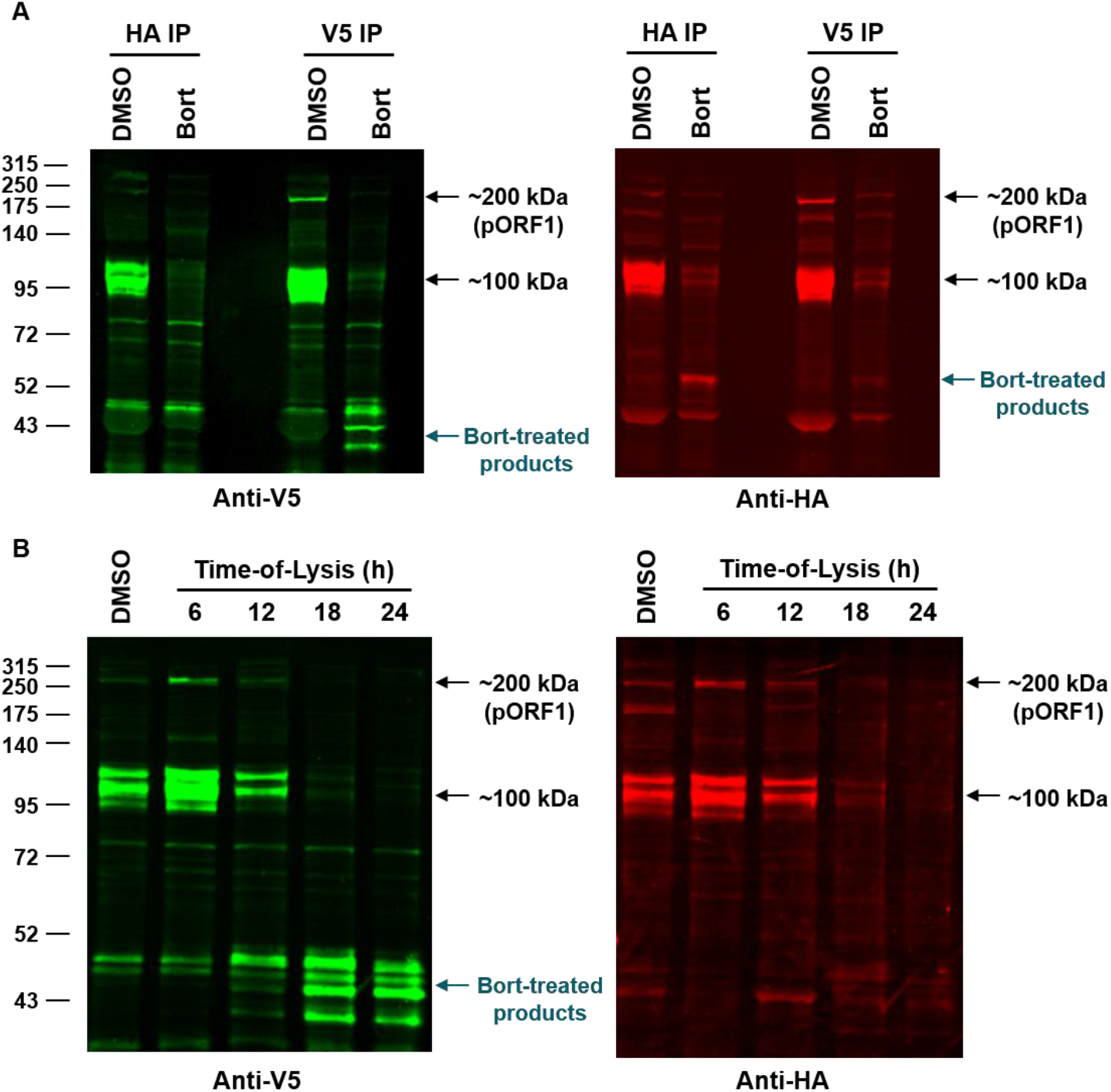
Bortezomib treatment disrupts HEV ORF1 proteolysis. (A) Huh7-SGR-G3-HA/V5 were treated with 1000 nM bortezomib or vehicle control (DMSO), with cells lysed at 24 h post-addition. Cell lysates were subject to reciprocal IPs, which were performed using anti-HA or anti-V5 nanobodies. This was followed by western blotting, probing for both anti-V5 and anti-HA. The native ORF1 products are indicated (∼200 and ∼100 kDa) alongside bortezomib treatment mediated products (blue arrows). (B) Time-course analysis of HEV ORF1 stability and replication under bortezomib treatment. Huh7-SGR-G3-HA/V5 were treated with 1000 nM bortezomib or DMSO. Cells were lysed at 6, 12, 18, and 24 h post-treatment, and subjected to IP using anti-V5 nanobodies. IP samples were subject to western blotting with anti-V5 or anti-HA antibodies.

Bortezomib treatment resulted in complete loss of the putative full-length ∼200 kDa and truncated/cleaved ∼100 kDa pORF1 proteins after 24 h compared to DMSO control in both IPs (Figure 7A). Alongside the loss of these pORF1 species, we also observed the appearance of additional lower molecular-weight proteins (∼45-55 kDa). After IP with anti-HA, a lower molecular weight product of ∼53 kDa was detected when probed with anti-HA, which was not visible when probed with anti-V5 (Figure 7A, merged blot shown in Figure S5A). In comparison, after IP with anti-V5, a distinct band of ∼45 kDa was observed when western blotted for anti-V5, yet these products were not observed when probed with anti-HA. Together, these data confirm previous observations suggesting that the proteasome is a key component required for HEV replication and pORF1 processing. However, we expand on these findings and show proteasome inhibition also generates lower molecular-weight pORF1 products, likely through alternative processing between the PCP and HVR domains.

To examine the kinetics and stability of pORF1, and the downstream effect on HEV replication, Huh7-SGR-G3-HA/V5 cells were treated with 1000 nM bortezomib and pORF1 expression examined at multiple timepoints. Lysates were harvested at 6, 12, 18, and 24 h post-bortezomib treatment, alongside a representative DMSO control. Each lysate was then subject to anti-V5 IP, followed by western blotting with either anti-V5 or anti-HA (Figure 7B, merged blot shown in Figure S5B).

This revealed that the uncleaved pORF1 (∼200 kDa) and ∼100 kDa products were only detectable at 6 and 12 h, with these substantially less abundant by 18 h, and completely absent by 24 h post-treatment (Figure 7B). From 12 h post-treatment onwards, the loss of pORF1 was accompanied by the appearance of lower molecular weight products (∼45 kDa) that remain until 24 h. To identify if these changes coincided with reduced HEV replication, Huh7 cells were electroporated with SGR-G3-nLuc expressing RNA and treated with 1000 nM bortezomib. At 6, 12, 18, and 24 h post-treatment, luciferase activity was examined, alongside a respective DMSO control (a schematic for this protocol is shown in Figure S5C). Initially, luciferase activity at 0 hours post-treatment (inhibitor added and the cells immediately lysed) was utilised as a control, and as expected no significant difference between DMSO and bortezomib-treated cells was observed. Interestingly, a significant decrease in nLuc expression was measured at all other time points examined (Figure S5D), suggesting that HEV replication is dependent on proteasome-associated ORF1 proteolytic cleavage.

Taken together, these findings suggest that the pORF1 polyprotein and/or the ∼100 kDa ORF1 cleavage products are essential for HEV replication. However, we hypothesise that proteasomal inhibition prevents HEV replication generating non-functional ∼53 and ∼45 kDa pORF1 products, which most likely represent pORF1 degradation/additional cleavage.

## Discussion

HEV has emerged as a significant cause of viral hepatitis, yet many fundamental aspects of its molecular biology and pathogenesis remain poorly defined. In particular, there is limited information surrounding the cellular proteins that interact with the viral non-structural polyprotein pORF1. To address this, we employed transposon-mediated mutagenesis to probe HEV for regions permissive to epitope insertion and generated a dual-tagged SGR for proteomic profiling of pORF1-associated factors.

Following a similar strategy to previous studies, we systematically screened pORF1 to identify regions permissive to epitope insertions without disrupting viral RNA replication (Szkolnicka et al., 2019; Metzger et al., 2022). Consistent with these previous studies, our transposon-mediated insertional screening also identified clusters of insertion sites within the Y-domain, HVR, and N-and C-terminus of the RdRp. Most insertions appeared to avoid regions of predicted protein secondary structure. These results also agree with studies that identified naturally occurring insertions, particularly in the HVR, that can modulate HEV replication (Shukla et al., 2011). When we tested a selection of insertion sites in a nano-luciferase expressing SGR many did not show luciferase expression above the non-replicating control. This observation likely reflects the relative sensitivities of the colony formation assay used for selection and the nano-luciferase assay used to measure transient replication. Alternatively, compensatory mutations or trans-complementation could have occurred during transposon selection that we were not able to identify in the post-selection data. However, uniquely in our dataset, we identified one insertion site within the PCP domain that was highly replication competent. Given that the PCP domain remains poorly characterised and its role during replication is largely unknown, we introduced an HA-epitope after residue 488 within the PCP domain. Our data suggest this PCP^HA^ insertion is permissive for replication in both G1 and G3 SGRs and compatible with the previously identified insertions in the HVR (Szkolnicka et al., 2019). This extends the toolkit available to study HEV by showing that both the PCP and HVR domains can accommodate epitope insertions without disrupting replication, enabling studies of pORF1 organisation and interdomain interactions.

To investigate protein-protein interactions between pORF11 and host factors important for replication, we used a stable cell line expressing PCP^HA^ (Huh7-SGR-G3-HA/V5). Following IP of lysates from Huh7-SGR-G3-HA/V5 cells, TMT-based LC-MS identified a distinct set of host factors not previously reported, with strong enrichment of proteasome-associated proteins, including seven protein subunits of the 19S regulatory complex of the 26S proteasome. These subunits function to recognise ubiquitinated targets, remove their ubiquitin chains, unfold substrate proteins, and deliver them to the 20S proteasome, the catalytic component of the 26S complex. (Glickman and Ciechanover, 2002). In agreement with previous studies (Karpe and Meng, 2012), proteasomal inhibition using bortezomib significantly reduced HEV replication across genotypes 1 and 3. However, our LC-MS dataset also uniquely identified several proteins associated with the ERAD pathway, which targets ubiquitin proteins to the 26S proteasome. Pharmacological inhibition of several ORF1-interacting ERAD proteins, each involved in different stages of the pathway, significantly inhibited HEV replication. Collectively, these findings extend previous studies and provide direct molecular evidence that pORF1 utilises the ERAD ubiquitin-proteasome system to support replication. This is also in agreement with a recent study implicating the proteasome in processing of pORF1 during replication (Zhang et al., 2025). These authors reported a ∼120 kDa pORF1 proteasomal-derived product that lacks the helicase and RdRp domains and modulates SMAD signalling. This product appears similar to the ∼100 kDa pORF1-derived products identified in our study.

Interestingly, treating Huh7-SGR-G3-HA/V5 cells with bortezomib for 24 h resulted in aberrant pORF1 processing, with the loss of native (∼200 kDa and ∼100 kDa proteins) pORF1 products and the concurrent appearance of multiple lower molecular weight proteins (∼40 kDa). The loss of native pORF1 product also coincided with a loss of HEV replication, suggesting these lower molecular-weight proteins cannot support replication. Together, these data are consistent with a model where pORF1 is trafficked through the ERAD pathway for proteasomal proteolysis to generate the ∼100-120 kDa pORF1 product to support replication. Inhibition of this process likely causes accumulation of pORF1 within the ER, where it undergoes aberrant proteolysis by ER-resident proteases. It is also possible that our observed ∼100 kDa pORF1 product is not the result of a cleavage event, but of alternative additional start codons, or early ribosomal release. Further work is ongoing to identify the nature of the lower molecular pORF1 products in our study.

Proteasomal degradation has also been shown to be important for the replication of other viruses, such as hepatitis C virus (HCV), by regulating of the non-structural proteins NS5A and NS5B. NS5A undergoes ubiquitin-dependent proteasomal degradation, a process enhanced by zinc mesoporphyrin (ZnMP), which suppresses viral RNA replication, while NS5B interacts with the ubiquitin-like protein hPLIC1, which links it to the proteasome and promotes polyubiquitylation and degradation to control its stability (Shoji, 2012). Beyond their degradative roles, 19S subunits are also recognised to support viral gene expression in other viruses such as HIV-1 and human cytomegalovirus, highlighting how viruses can exploit these components for replication (Lassot et al., 2007; Winkler and Kalejta, 2014). The identification of 19S components interacting with HEV pORF1 suggests that HEV may similarly exploit the 19S regulatory particle. Further clarification is needed to understand HEV’s reliance on an active 26S proteasome for replication and the potential repurposing of FDA-approved compounds such as bortezomib to inhibit HEV replication.

From our IP data, we were able to detect both ∼100-120 kDa pORF1 products as well as full-length pORF1. It is unclear whether all pORF1 is trafficked through the proteasome and the full-length pORF1 observed is part of a dynamic equilibrium, or whether a fraction of pORF1 remains unprocessed. The ∼100 kDa pORF1-proteasomal product was reported to lack the helicase and RdRp domains; therefore, it cannot support RNA replication alone. Therefore, if proteasomal proteolysis is required to directly support viral RNA replication it could potentially be through release of the C-terminal RdRp-containing fragment. Alternatively, the 120 kDa product could also provide role(s) required for replication alongside SMAD signalling. Studies on HEV pORF1 processing have yielded conflicting results across multiple experimental systems, including multiple products identified and proteases implicated (Ropp et al., 2000; Sehgal et al., 2006; Kanade et al., 2019; Metzger et al., 2022; Pierce et al., 2023) including the suggestion that pORF1 functions as an unprocessed unit (Ansari et al., 2000; Suppiah et al., 2011; Perttilä et al., 2013; Ding et al., 2018; Szkolnicka et al., 2019; Ju et al., 2020).

Interestingly, immunofluorescence analysis of the Huh7-SGR-G3-HA/V5 cell line revealed a predominantly cytoplasmic distribution with discrete punctate foci of both PCP and HVR, often enriched in the perinuclear region, consistent with prior studies using epitope-labelled HVR (Perttilä et al., 2013; Szkolnicka et al., 2019; Metzger et al., 2022). However, epitope tags in two pORF1 domains allows us to show that PCP puncta overlap with HVR-positive structures, consistent with our IP data and prior observations (Metzger et al., 2022). Co-immunofluorescent analysis of Huh7-SGR-G3-HA/V5 for dsRNA, a hallmark of positive-sense viral RNA replication (Weber et al., 2006), demonstrated that most HVR co-localised with dsRNA (∼60%). However, the majority of dsRNA puncta observed within Huh7-SGR-G3-HA/V5 did not co-localise with HVR (∼80%). These observations are consistent with only a proportion of pORF1 localising to sites of viral replication and the existence of different pools of pORF1 after alternative polyprotein processing, potentially from different proteases. An alternative explanation for this is that persistent HEV replication produces relatively stable dsRNA intermediates that can persist independently of pORF1 components. Similar observations have been observed for cells persistently infected with porcine reproductive and respiratory syndrome virus (PRRSV) (Guo et al., 2018). Here, the author’s report that the persistently PRRSV-infected cells had a high level of dsRNA, which had less co-localisation with the examined viral proteins, when compared to acute infection.

In conclusion, our data point to key roles for the proteasome and the ERAD pathway in HEV genome replication. Future work will focus on determining the mechanistic details of this requirement. Key to this will be identifying and characterising of the different pORF1 proteolytic cleavage products observed in the presence and absence of proteasome or ERAD inhibitors.

## Materials and Methods

### Cell lines and plasmids

Huh7 cells (ATCC), were maintained in Dulbecco’s modified Eagle’s medium (DMEM) with glutamine (Sigma-Aldrich) supplemented with 10% (v/v) foetal bovine serum, 1% (v/v) non-essential amino acids, 10 mM HEPES, 50 U/mL penicillin and 50 μg/mL streptomycin. The plasmid containing the G1 HEV (Sar55 strain; GenBank accession no. AF444002) SGR expressing nano-luciferase (pSK-E2-nLuc) has been previously described (Herod et al., 2022). The plasmid containing the G3 HEV (G3-HEV83-2-27 strain; GenBank accession no AB740232) SGR expressing nano-luciferase (pUC-HEV83-2) has been described previously (Shiota et al., 2013). Negative control pSK-E2-nLuc and pUC-HEV-83-2 SGRs were generated containing a double point mutation in the RdRp active site GDD motif (GNN), as previously described (Pierce et al., 2023). Mutations within these plasmids were performed by standard two-step overlapping PCR mutagenesis. For G1 HEV SGR-expressing nano-luciferase (nLuc) transposon sites, insertion PCRs were performed using Q5^®^ High-Fidelity 2X Master Mix (NEB), according to the manufacturer’s instructions.

### *In vitro* transcription

The G1 and G3 HEV replicon plasmids were linearised with *Bgl*II (NEB) or *Hin*dIII (NEB), respectively, before generating T7 *in vitro* transcribed RNA as previously described (Pierce et al., 2023). RNA was purified using an RNA clean and concentrate kit (Zymo Research) and the quality was checked using MOPS/formaldehyde agarose gel electrophoresis.

### Transposon mutagenesis

pUC-SK-E2-ORF1 plasmid (150 ng) was used for *in vitro* transposon mutagenesis (Tn) carried out using the Mutation Generation System Kit (Thermo Scientific) according to the manufacturer’s instructions. Tn was halted by heating the reaction to 95 °C for 10 mins. The resulting Tn library was electroporated into Electro-Ten-Blue cells (Agilent) (electrocompetent E. coli) in 0.1 mm gap cuvettes under the following conditions: 250 V, 22.5 kV/cm field strength, 200 Ω resistance and 25 μF capacitance, prior to immediate recovery in Super Optimal Broth with Catabolic repression (SOC) (Sigma-Aldrich) preheated to 36°C. Electroporated cells were subsequently spread onto LB agar plates containing ampicillin (100 μg/mL) and chloramphenicol (10 μg/mL). Plasmid DNA was extracted from pooled colonies using a Monarch plasmid miniprep kit (NEB). The ORF1 coding region from the library was removed via restriction digestion and subsequently cloned into pSK-E2-Neo-MfeI. This plasmid was digested to remove the chloramphenicol resistance cassette, and the resulting re-ligated plasmid formed the library plasmid pSK-E2-Neo-Tn-lib.

### Sequencing & Bioinformatic analysis

The pre-selected transposon library plasmid (pSK-E2-Neo-Tn-Lib) was whole-plasmid sequenced by Microbes NG (UK) on an Illumina platform with 60X coverage. Post-selected library amplicons were also sequenced via paired-end reads by Microbe NG, on an Illumina platform with 150X coverage. Sequencing reads from pre-selection (“Pre”) and post-selection (“Post”) libraries were processed to identify transposon–virus junctions in line with previous literature (Heaton et al., 2013; Fulton et al., 2018). Read pairs were screened using cutadapt (v.5.2) with the parameters: -g TGCGGCCGCA –overlap 10 -e 0.05 –discard-untrimmed -m 20, to identify the presence of a fixed 10-nucleotide transposon tag (TGCGGCCGCA), requiring a full-length match with ≤ 5% mismatches (Martin, 2011). The tag and all upstream sequence were removed and only read pairs in which read 1 contained the tag and remained ≥ 20-nt after trimming were retained. In addition, all reads were mapped to the HEV genome to determine coverage of the whole genome and to normalise to insertion site per million. Trimmed paired-end reads were aligned to the hepatitis E virus reference genome (AF444002.1) using Bowtie2 (v.2.5.4) (Langmead and Salzberg, 2012) in very-sensitive mode, and only high-confidence alignments (mapping quality ≥ 20, no soft clipping) were used for insertion calling, resulting bam files were sorted and indexed with samtools (v.1.15.1). Insertion coordinates extracted using samtools and awk (v.5.3.1) commands; and defined as the nucleotide immediately adjacent to the first mapped viral base of each trimmed read, adjusted for strand orientation, and identical sites were collapsed to generate per-site insertion counts for each library. In R, insertion sites were aggregated into non-overlapping 10-nucleotide bins, and the log2 frequency was visualised for both “Pre” and “Post” libraries where the number of reads per bin >1; a dashed line highlights 3 reads per bin.

### Generation of Huh7-SGR-G3-HA/V5

To establish the Huh7-SGR-G3-HA/V5 (PCP^HA^ and HVR^V5^) cells Huh7 were electroporated with the SGR-expressing NPTII. Briefly, cells were seeded into 6-well plates at 10^6^, 10^5^ and 10^4^ cells per well, with the addition of untransfected Huh7 cells to achieve a total of 2×10^6^ cells per well. After 24 h, cells expressing NPTII were selected by addition of 500 µg/ml geneticin (G418 Sulfate, Gibco Invitrogen) with daily monitoring of cell death and confluency. To sustain optimal selection, G418 was adjusted between 250 µg/mL and 750 µg/mL. Cells were expanded following selection, with large colonies present after 2 weeks.

### Replication assays

Replicon experiments were conducted as previously described (Herod et al., 2022). Briefly, Huh7 cells were detached by trypsin, washed twice in ice-cold DEPC-treated PBS and re-suspended at 1 × 10^7^ cells/mL in DEPC-treated PBS. Subsequently 400 μL of cells were mixed with 2 μg of RNA transcript, transferred to a 4 mm gap electroporation cuvette (SLS, UK) and pulsed at 260 V, 25 ms pulse length in a Bio-Rad Gene Pulser (Bio-Rad). Electroporated cells were recovered into 4 mL media, seeded into replicate 6-well tissue culture vessels, and replication measured at 24 h intervals using a Nano-Glo luciferase assay system (Promega). For inhibitor treatment the electroporated cells were seeded into replicate 12-well plates, allowed to adhere for 24 h before the media was replaced with fresh media containing bortezomib (Stratech Scientific Ltd), CB-5083 (TargetMol Chemical Inc), TAK-243 (TargetMol Chemical Inc) and HA15 (TargetMol Chemical Inc). Cell supernatants were harvested at 24 and 48 h post-addition of inhibitor, and luciferase activity was measured. The cell viability experiments were conducted by seeding cells into 96-well plates, allowing them to adhere for 24 h before measurement of cell viability at 48/72 h later using the CellTiter AQueous One solution (Promega, USA), following manufacturer’s instructions.

### Immunofluorescence

Huh7-SGR-G3-HA/V5 cells were seeded at 1 × 10^5^ cells/well, followed by incubation at 37°C. After 24 h, cells were washed two times in PBS prior to fixation in 4% paraformaldehyde for 15 min at room temperature. After fixation, the cells were washed three times in PBS and then incubated in permeabilisation buffer (0.3% Triton X-100, 1% bovine serum albumin (BSA) in PBS) for 10 min at room temperature. Following permeabilisation, the monolayers were washed three times with PBS and incubated with blocking buffer (1% BSA in PBS) for 1 h. Subsequently, primary antibody was diluted in the same blocking buffer. Rabbit anti-V5 (1:500 EPR12989; Abcam) and Mouse anti-HA (1:500 H3663; Sigma-Aldrich), detected with AlexaFluor secondary antibodies. Coverslips were washed a final time in PBS before mounting in ProLong Glass with NucBlue (Thermo Fisher Scientific, USA). Images were captured using a Zeiss LSM-880 confocal microscope. Mander’s Overlap Co-efficient was calculated using the JACoP function in ImageJ on ten separate images, for which mean and standard deviation values were attributed.

### Co-Immunoprecipitation

Huh7-SGR-G3-HA/V5 were cultured in T175 flasks and lysed, with rotation, on ice for 30 min in 1 mL of Pierce™ IP Lysis Buffer (Thermo Fisher Scientific) supplemented with Halt™ Protease Inhibitor Cocktail (Thermo Fisher Scientific). Lysates were cleared by centrifugation at 17,000 x g for 15 minutes at 4°C, and a 50 μL aliquot taken for input control. Two separate co-immunoprecipitation reactions were performed. Anti-HA pulldown was carried out using 200 μL of Pierce™ Anti-HA Magnetic Beads (Thermo Fisher Scientific), washed once with PBS containing 0.02% Tween-20 (PBS-T), then incubated with cleared lysate overnight at 4°C with gentle agitation. Beads were washed three times with 800 μL of PBS-T. During the final wash step, beads were transferred to fresh tubes to minimise non-specific background. After washing, beads were resuspended in 50 μL of PBS-T (0.02%) and stored at −80°C prior to TMT-MS analysis. V5-tagged proteins were immunoprecipitated using 200 μL of ChromoTek V5-Trap® Magnetic Particles M-270 (Proteintech), processed according to the manufacturer’s protocol and incubated under identical conditions.

### Western blotting

Western blot samples were prepared by adding of 4× Laemmli sample buffer containing 10% v/v β-mercaptoethanol (prepared by mixing 4 μL sample buffer with 12 μL 0.02% PBS-T), followed by boiling at 95°C for 5 min. Proteins were resolved using an 8% SDS-PAGE gel and transferred to PVDF membranes. Membranes were blocked in Licor blocking buffer 1:1 with 0.1% PBS-T. Primary antibodies were then added mouse anti-GAPDH (sc-32233, Santa Cruz Biotechnology), 1:5000, mouse anti-HA (H3663, Sigma-Aldrich), 1:1000, rabbit anti-V5 (SY30-01, Invitrogen), 1:1000, mouse anti-neomycin phosphotransferase II (4B4D1, Invitrogen), 1:2000. Blots were then subsequently incubated with secondary antibodies goat anti-rabbit (DyLight™ 800) and goat anti-mouse (DyLight™ 680), prior to Licor fluorescent imaging.

### TMT Labelling and High pH reversed-phase chromatography

Immuno-isolated samples were reduced (10mM TCEP, 55°C for 1 h), alkylated (18.75mM iodoacetamide, room temperature for 30 min.) and then digested from the beads with trypsin (1.25µg trypsin; 37°C, overnight). The resulting peptides were then labelled with Tandem Mass Tag (TMTpro) sixteen-plex reagents according to the manufacturer’s protocol (Thermo Fisher Scientific, Loughborough, LE11 5RG, UK) and the labelled samples were pooled and desalted using a SepPak cartridge according to the manufacturer’s instructions (Waters, Milford, Massachusetts, USA). Eluate from the SepPak cartridge was evaporated to dryness and resuspended in buffer A (20 mM ammonium hydroxide, pH 10) prior to fractionation by high pH reversed-phase chromatography using an Ultimate 3000 liquid chromatography system (Thermo Scientific). In brief, the sample was loaded onto an XBridge BEH C18 Column (130Å, 3.5 µm, 2.1 mm X 150 mm, Waters, UK) in buffer A and peptides eluted with an increasing gradient of buffer B (20 mM Ammonium Hydroxide in acetonitrile, pH 10) from 0-95% over 60 minutes. The resulting fractions (concatenated into 4 in total) were evaporated to dryness and resuspended in 1% formic acid prior to analysis by nano-LC MSMS using an Orbitrap Fusion Tribrid mass spectrometer (Thermo Scientific).

### Nano-LC Mass Spectrometry

High pH RP fractions were further fractionated using an Ultimate 3000 nano-LC system in line with an Orbitrap Fusion Tribrid mass spectrometer (Thermo Scientific). In brief, peptides in 1% (vol/vol) formic acid were injected onto an Acclaim PepMap C18 nano-trap column (Thermo Scientific). After washing with 0.5% (vol/vol) acetonitrile 0.1% (vol/vol) formic acid peptides were resolved on a 250 mm × 75 μm Acclaim PepMap C18 reverse phase analytical column (Thermo Scientific) over a 150 min organic gradient, using 7 gradient segments (1-6% solvent B over 1min., 6-15% B over 58min., 15-32%B over 58min., 32-40%B over 5min., 40-90%B over 1min., held at 90%B for 6min and then reduced to 1%B over 1min.) with a flow rate of 300 nl min−1. Solvent A was 0.1% formic acid and Solvent B was aqueous 80% acetonitrile in 0.1% formic acid. Peptides were ionised by nano-electrospray ionisation at 2.0kV using a stainless-steel emitter with an internal diameter of 30 μm (Thermo Scientific) and a capillary temperature of 275°C.

All spectra were acquired using an Orbitrap Fusion Tribrid mass spectrometer controlled by Xcalibur 2.1 software (Thermo Scientific) and operated in data-dependent acquisition mode using an SPS-MS3 workflow. FTMS1 spectra were collected at a resolution of 120 000, with an automatic gain control (AGC) target of 400 000 and a max injection time of 100ms. Precursors were filtered with an intensity range from 5000 to 1E20, according to charge state (to include charge states 2-6) and with monoisotopic peak determination set to peptide. Previously interrogated precursors were excluded using a dynamic window (60s +/−10ppm). The MS2 precursors were isolated with a quadrupole isolation window of 1.2m/z. ITMS2 spectra were collected with an AGC target of 10 000, a max injection time of 70 ms and a CID collision energy of 35%.

For FTMS3 analysis, the Orbitrap was operated at 30 000 resolution with an AGC target of 50 000 and a max injection time of 105ms. Precursors were fragmented by high energy collision dissociation (HCD) at a normalised collision energy of 55% to ensure maximal TMT reporter ion yield. Synchronous Precursor Selection (SPS) was enabled to include up to 10 MS2 fragment ions in the FTMS3 scan.

### Data Analysis

The raw data files were processed and quantified using Proteome Discoverer software v2.4 (Thermo Scientific) and searched against the UniProt Human database (downloaded January 2025; 83095 sequences) and the HEV_83-2-HA-V5 tag ORF1 sequence using the SEQUEST HT algorithm. Peptide precursor mass tolerance was set to 10 ppm, and MS/MS tolerance was set to 0.6Da. Search criteria included oxidation of methionine (+15.995Da), acetylation of the protein N-terminus (+42.011Da), methionine loss form the protein N-terminus (−131.04Da), methionine loss plus acetylation of the protein N-terminus (−89.03Da), carbamidomethylation of cysteine (+57.021Da) and palmitoylation of cysteine (+238.23Da) as variable modifications and addition of the TMTpro mass tag (+304.207Da) to peptide N-termini and lysine as fixed modifications. Searches were performed with full tryptic digestion and a maximum of 2 missed cleavages were allowed. The reverse database search option was enabled, and all data were filtered to satisfy a false discovery rate (FDR) of 5%. The data generated were assessed based on the enrichment of proteins isolated from the Huh7-SGR-G3-HA/V5 samples when compared to Huh7. Statistical analysis, using an unpaired T test, was performed to increase confidence in identifying significantly enriched genes. Subsequently, genes that met the significance criteria (false discovery rate FDR-pass, p-value <0.05 and log2FC >1) were selected for subsequent downstream functional enrichment analysis. GO enrichment analysis based on Cytoscape software with ClueGO was used to connect proteins linked via cellular pathways and GO molecular function.

## Acknowledgements

We thank Patrizia Farci (National Institute of Allergy and Infectious Diseases, Bethesda, USA) for the genotype 1 HEV SGR and Koji Ishii (National Institute of Infectious Disease, Tokyo, Japan) for the genotype 3 HEV SGR.

## Author contributions

Conceptualization: KMA, FJTB, OB, MH and MRH.

Investigation: KMA, FJTB, OB, EMB, CEW, MK and RPR.

Supervision: MH and MRH.

Writing – original draft: KMA, FJTB, OB, RPR, MH and MRH.

Writing – review and editing: KMA, FJTB, OB, EMB, CEW, MK, RPR, MH and MRH.

## Data availability

The authors declare that the main data supporting the findings of this study are available within the article and its supplemental material. The mass spectrometry proteomics data have been deposited to the ProteomeXchange Consortium via the PRIDE partner repository with the data set identifier PXD084724.

## References

Adlhoch, C., Avellon, A., Baylis, S.A., Ciccaglione, A.R., Couturier, E., De Sousa, R., Epštein, J., Ethelberg, S., Faber, M., Fehér, Á., Ijaz, S., Lange, H., Manďáková, Z., Mellou, K., Mozalevskis, A., Rimhanen-Finne, R., Rizzi, V., Said, B., Sundqvist, L., Thornton, L., Tosti, M.E., Van Pelt, W., Aspinall, E., Domanovic, D., Severi, E., Takkinen, J. and Dalton, H.R. 2016. Hepatitis E virus: Assessment of the epidemiological situation in humans in Europe, 2014/15. Journal of Clinical Virology. 82, pp.9–16.

Ansari, I.H., Nanda, S.K., Durgapal, H., Agrawal, S., Mohanty, S.K., Gupta, D., Jameel, S. and Panda, S.K. 2000. Cloning, sequencing, and expression of the hepatitis E virus (HEV) nonstructural open reading frame 1 (ORF1). Journal of Medical Virology. 60(3), pp.275–283.

Arakawa, Y., Jo, U., Kumar, S., Sun, N.-Y., Elloumi, F., Thomas, A., Roper, N., Varghese, D.G., Takebe, N., Zhang, X., Ceribelli, M., Holland, D.O., Beck, E., Itkin, Z., McKnight, C., Wilson, K.M., Travers, J., Klumpp-Thomas, C., Thomas, C.J., Hoang, C.D., Hernandez, J.M., Del Rivero, J. and Pommier, Y. 2024. Activity of the Ubiquitin-activating Enzyme Inhibitor TAK-243 in Adrenocortical Carcinoma Cell Lines, Patient-derived Organoids, and Murine Xenografts. Cancer Research Communications. 4(3), pp.834–848.

Bigna, J.J., Modiyinji, A.F., Nansseu, J.R., Amougou, M.A., Nola, M., Kenmoe, S., Temfack, E. and Njouom, R. 2020. Burden of hepatitis E virus infection in pregnancy and maternofoetal outcomes: a systematic review and meta-analysis. BMC Pregnancy and Childbirth. 20(1), p.426.

Boccadoro, M., Morgan, G. and Cavenagh, J. 2005. Preclinical evaluation of the proteasome inhibitor bortezomib in cancer therapy. Cancer Cell International. 5(1), p.18.

Buchanan, F.J.T., Chen, S., Harris, M. and Herod, M.R. 2023. The hepatitis E virus ORF1 hypervariable region confers partial cyclophilin dependency. Journal of General Virology. 104(11).

Chen, D., Frezza, M., Schmitt, S., Kanwar, J. and P. Dou, Q. 2011. Bortezomib as the First Proteasome Inhibitor Anticancer Drug: Current Status and Future Perspectives. Current Cancer Drug Targets. 11(3), pp.239–253.

Dalton, H.R., Bendall, R.P., Keane, F.E., Tedder, R.S. and Ijaz, S. 2009. Persistent Carriage of Hepatitis E Virus in Patients with HIV Infection. New England Journal of Medicine. 361(10), pp.1025–1027.

Debing, Y., Gisa, A., Dallmeier, K., Pischke, S., Bremer, B., Manns, M., Wedemeyer, H., Suneetha, P.V. and Neyts, J. 2014. A Mutation in the Hepatitis E Virus RNA Polymerase Promotes Its Replication and Associates With Ribavirin Treatment Failure in Organ Transplant Recipients. Gastroenterology. 147(5), pp.1008–1011.e7.

Ding, Q., Heller, B., Capuccino, J.M.V., Song, B., Nimgaonkar, I., Hrebikova, G., Contreras, J.E. and Ploss, A. 2017. Hepatitis E virus ORF3 is a functional ion channel required for release of infectious particles. Proceedings of the National Academy of Sciences. 114(5), pp.1147–1152.

Ding, Q., Nimgaonkar, I., Archer, N.F., Bram, Y., Heller, B., Schwartz, R.E. and Ploss, A. 2018. Identification of the Intragenomic Promoter Controlling Hepatitis E Virus Subgenomic RNA Transcription X.-J. Meng, ed. mBio. 9(3), pp.e00769–18.

Eletto, Davide, Eletto, Daniela, Dersh, D., Gidalevitz, T. and Argon, Y. 2014. Protein Disulfide Isomerase A6 Controls the Decay of IRE1α Signaling via Disulfide-Dependent Association. Molecular Cell. 53(4), pp.562–576.

Fulton, B.O., Sun, W., Heaton, N.S. and Palese, P. 2018. The Influenza B Virus Hemagglutinin Head Domain Is Less Tolerant to Transposon Mutagenesis than That of the Influenza A Virus T. S. Dermody, ed. Journal of Virology. 92(16), pp.e00754–18.

Glickman, M.H. and Ciechanover, A. 2002. The Ubiquitin-Proteasome Proteolytic Pathway: Destruction for the Sake of Construction. Physiological Reviews. 82(2), pp.373–428.

Guo, R., Shang, P., Carrillo, C.A., Sun, Z., Lakshmanappa, Y.S., Yan, X., Renukaradhya, G.J., McGill, J., Jaing, C.J., Niederwerder, M.C., Rowland, R.R.R. and Fang, Y. 2018. Double-stranded viral RNA persists in vitro and in vivo during prolonged infection of porcine reproductive and respiratory syndrome virus. Virology. 524, pp.78–89.

Haase, M. and Fitze, G. 2016. HSP90AB1: Helping the good and the bad. Gene. 575(2), pp.171–186.

Heaton, N.S., Sachs, D., Chen, C.-J., Hai, R. and Palese, P. 2013. Genome-wide mutagenesis of influenza virus reveals unique plasticity of the hemagglutinin and NS1 proteins. Proceedings of the National Academy of Sciences. 110(50), pp.20248–20253.

Herod, M.R., Loundras, E.-A., Ward, J.C., Tulloch, F., Rowlands, D.J. and Stonehouse, N.J. 2015. Employing transposon mutagenesis to investigate foot-and-mouth disease virus replication. Journal of General Virology. 96(12), pp.3507–3518.

Herod, M.R., Ward, J.C., Tuplin, A., Harris, M., Stonehouse, N.J. and McCormick, C.J. 2022. Positive strand RNA viruses differ in the constraints they place on the folding of their negative strand. RNA. 28(10), pp.1359–1376.

Ju, X., Xiang, G., Gong, M., Yang, R., Qin, J., Li, Y., Nan, Y., Yang, Y., Zhang, Q.C. and Ding, Q. 2020. Identification of functional cis-acting RNA elements in the hepatitis E virus genome required for viral replication Z. Feng, ed. PLOS Pathogens. 16(5), p.e1008488.

Ju, X., Yu, Y., Ren, W., Dong, L., Meng, X., Deng, H., Nan, Y. and Ding, Q. 2023. The PRMT5/WDR77 complex restricts hepatitis E virus replication S. M. Best, ed. PLOS Pathogens. 19(6), p.e1011434.

Kamar, N., Garrouste, C., Haagsma, E.B., Garrigue, V., Pischke, S., Chauvet, C., Dumortier, J., Cannesson, A., Cassuto-Viguier, E., Thervet, E., Conti, F., Lebray, P., Dalton, H.R., Santella, R., Kanaan, N., Essig, M., Mousson, C., Radenne, S., Roque-Afonso, A.M., Izopet, J. and Rostaing, L. 2011. Factors Associated With Chronic Hepatitis in Patients With Hepatitis E Virus Infection Who Have Received Solid Organ Transplants. Gastroenterology. 140(5), pp.1481–1489.

Kanade, G.D., Pingale, K.D. and Karpe, Y.A. 2019. Protein Interactions Network of Hepatitis E Virus RNA and Polymerase With Host Proteins. Frontiers in Microbiology. 10, p.2501.

Karpe, Y.A. and Meng, X.-J. 2012. Hepatitis E Virus Replication Requires an Active Ubiquitin-Proteasome System. Journal of Virology. 86(10), pp.5948–5952.

Kisselev, A.F. and Goldberg, A.L. 2001. Proteasome inhibitors: from research tools to drug candidates. Chemistry & Biology. 8(8), pp.739–758.

Koonin, E.V., Gorbalenya, A.E., Purdy, M.A., Rozanov, M.N., Reyes, G.R. and Bradley, D.W. 1992. Computer-assisted assignment of functional domains in the nonstructural polyprotein of hepatitis E virus: delineation of an additional group of positive-strand RNA plant and animal viruses. Proceedings of the National Academy of Sciences. 89(17), pp.8259–8263.

Langmead, B. and Salzberg, S.L. 2012. Fast gapped-read alignment with Bowtie 2. Nature Methods. 9(4), pp.357–359.

Lassot, I., Latreille, D., Rousset, E., Sourisseau, M., Linares, L.K., Chable-Bessia, C., Coux, O., Benkirane, M. and Kiernan, R.E. 2007. The Proteasome Regulates HIV-1 Transcription by Both Proteolytic and Nonproteolytic Mechanisms. Molecular Cell. 25(3), pp.369–383.

Le Moigne, R., Aftab, B.T., Djakovic, S., Dhimolea, E., Valle, E., Murnane, M., King, E.M., Soriano, F., Menon, M.-K., Wu, Z.Y., Wong, S.T., Lee, G.J., Yao, B., Wiita, A.P., Lam, C., Rice, J., Wang, J., Chesi, M., Bergsagel, P.L., Kraus, M., Driessen, C., Kiss Von Soly, S., Yakes, F.M., Wustrow, D., Shawver, L., Zhou, H.-J., Martin, T.G., Wolf, J.L., Mitsiades, C.S., Anderson, D.J. and Rolfe, M. 2017. The p97 Inhibitor CB-5083 Is a Unique Disrupter of Protein Homeostasis in Models of Multiple Myeloma. Molecular Cancer Therapeutics. 16(11), pp.2375–2386.

Li, P., Liu, J., Li, Y., Su, J., Ma, Z., Bramer, W.M., Cao, W., De Man, R.A., Peppelenbosch, M.P. and Pan, Q. 2020. The global epidemiology of hepatitis E virus infection: A systematic review and meta-analysis. Liver International. 40(7), pp.1516–1528.

Liu, Z., Liu, G., Ha, D.P., Wang, J., Xiong, M. and Lee, A.S. 2023. ER chaperone GRP78/BiP translocates to the nucleus under stress and acts as a transcriptional regulator. Proceedings of the National Academy of Sciences. 120(31), p.e2303448120.

Martin, M. 2011. Cutadapt removes adapter sequences from high-throughput sequencing reads. EMBnet.journal. 17(1), p.10.

Meng, X.-J. 2016. Expanding Host Range and Cross-Species Infection of Hepatitis E Virus R. C. Condit, ed. PLOS Pathogens. 12(8), p.e1005695.

Meng, X.-J. 2013. Zoonotic and Foodborne Transmission of Hepatitis E Virus. Seminars in Liver Disease. 33(01), pp.041–049.

Metzger, K., Bentaleb, C., Hervouet, K., Alexandre, V., Montpellier, C., Saliou, J.-M., Ferrié, M., Camuzet, C., Rouillé, Y., Lecoeur, C., Dubuisson, J., Cocquerel, L. and Aliouat-Denis, C.-M. 2022. Processing and Subcellular Localization of the Hepatitis E Virus Replicase: Identification of Candidate Viral Factories. Frontiers in Microbiology. 13, p.828636.

Mi, Z. and Graham, S.H. 2023. Role of UCHL1 in the pathogenesis of neurodegenerative diseases and brain injury. Ageing Research Reviews. 86, p.101856.

Montpellier, C., Wychowski, C., Sayed, I.M., Meunier, J.-C., Saliou, J.-M., Ankavay, M., Bull, A., Pillez, A., Abravanel, F., Helle, F., Brochot, E., Drobecq, H., Farhat, R., Aliouat-Denis, C.-M., Haddad, J.G., Izopet, J., Meuleman, P., Goffard, A., Dubuisson, J. and Cocquerel, L. 2018. Hepatitis E Virus Lifecycle and Identification of 3 Forms of the ORF2 Capsid Protein. Gastroenterology. 154(1), pp.211–223.e8.

Nimgaonkar, I., Ding, Q., Schwartz, R.E. and Ploss, A. 2018. Hepatitis E virus: advances and challenges. Nature Reviews Gastroenterology & Hepatology. 15(2), pp.96–110.

Perry, J.W., Chen, Y., Speliotes, E. and Tai, A.W. 2018. Functional Analysis of the Dengue Virus Genome Using an Insertional Mutagenesis Screen M. S. Diamond, ed. Journal of Virology. 92(7), pp.e02085–17.

Perttilä, J., Spuul, P. and Ahola, T. 2013. Early secretory pathway localization and lack of processing for hepatitis E virus replication protein pORF1. Journal of General Virology. 94(4), pp.807–816.

Pierce, D.M., Buchanan, F.J.T., Macrae, F.L., Mills, J.T., Cox, A., Abualsaoud, K.M., Ward, J.C., Ariëns, R.A.S., Harris, M., Stonehouse, N.J. and Herod, M.R. 2023. Thrombin cleavage of the hepatitis E virus polyprotein at multiple conserved locations is required for genome replication G. A. Belov, ed. PLOS Pathogens. 19(7), p.e1011529.

Purdy, M.A., Drexler, J.F., Meng, X.-J., Norder, H., Okamoto, H., Van Der Poel, W.H.M., Reuter, G., De Souza, W.M., Ulrich, R.G. and Smith, D.B. 2022. ICTV Virus Taxonomy Profile: Hepeviridae 2022: This article is part of the ICTV Virus Taxonomy Profiles collection. Journal of General Virology. 103(9).

Rehati, A., Abuduaini, B., Liang, Z., Chen, D. and He, F. 2023. Identification of heat shock protein family A member 5 (HSPA5) targets involved in nonalcoholic fatty liver disease. Genes & Immunity. 24(3), pp.124–129.

Ropp, S.L., Tam, A.W., Beames, B., Purdy, M. and Frey, T.K. 2000. Expression of the hepatitis E virus ORF1. Archives of Virology. 145(7), pp.1321–1337.

Saibil, H. 2013. Chaperone machines for protein folding, unfolding and disaggregation. Nature Reviews Molecular Cell Biology. 14(10), pp.630–642.

Sehgal, D., Thomas, S., Chakraborty, M. and Jameel, S. 2006. Expression and processing of the Hepatitis E virus ORF1 nonstructural polyprotein. Virology Journal. 3(1), p.38.

Shiota, T., Li, T.-C., Yoshizaki, S., Kato, T., Wakita, T. and Ishii, K. 2013. The Hepatitis E Virus Capsid C-Terminal Region Is Essential for the Viral Life Cycle: Implication for Viral Genome Encapsidation and Particle Stabilization. Journal of Virology. 87(10), pp.6031–6036.

Shoji, I. 2012. Roles of the two distinct proteasome pathways in hepatitis C virus infection. World Journal of Virology. 1(2), p.44.

Shukla, P., Nguyen, H.T., Torian, U., Engle, R.E., Faulk, K., Dalton, H.R., Bendall, R.P., Keane, F.E., Purcell, R.H. and Emerson, S.U. 2011. Cross-species infections of cultured cells by hepatitis E virus and discovery of an infectious virus–host recombinant. Proceedings of the National Academy of Sciences. 108(6), pp.2438–2443.

Subramani, C., Nair, V.P., Anang, S., Mandal, S.D., Pareek, M., Kaushik, N., Srivastava, A., Saha, S., Shalimar, Nayak, B., Ranjith-Kumar, C.T. and Surjit, M. 2018. Host-Virus Protein Interaction Network Reveals the Involvement of Multiple Host Processes in the Life Cycle of Hepatitis E Virus D. M. Knipe, ed. mSystems. 3(1), pp.e00135–17.

Suppiah, S., Zhou, Y. and Frey, T.K. 2011. Lack of Processing of the Expressed ORF1 Gene Product of Hepatitis E Virus. Virology Journal. 8(1), p.245.

Szkolnicka, D., Pollán, A., Da Silva, N., Oechslin, N., Gouttenoire, J. and Moradpour, D. 2019. Recombinant Hepatitis E Viruses Harboring Tags in the ORF1 Protein J.-H. J. Ou, ed. Journal of Virology. 93(19), pp.e00459–19.

Teterina, N.L., Lauber, C., Jensen, K.S., Levenson, E.A., Gorbalenya, A.E. and Ehrenfeld, E. 2011. Identification of tolerated insertion sites in poliovirus non-structural proteins. Virology. 409(1), pp.1–11.

Tian, G., Hu, C., Yun, Y., Yang, W., Dubiel, W., Cheng, Y. and Wolf, D.A. 2021. Dual roles of HSP70 chaperone HSPA1 in quality control of nascent and newly synthesized proteins. The EMBO Journal. 40(13), p.EMBJ2020106183.

Vembar, S.S. and Brodsky, J.L. 2008. One step at a time: endoplasmic reticulum-associated degradation. Nature Reviews Molecular Cell Biology. 9(12), pp.944–957.

Wang, Zhen, Guo, Z., Cao, C., Wang, Ziying, Huang, Z., Zhang, X., Zheng, Y., She, D., Zhu, H., Yu, L., Zhao, X., Xiang, D., Liao, S., He, X., Ji, X., Wu, C., Xu, C. and Xu, M. 2026. HSP90AB1-Mediated Ubiquitin-Proteasome Degradation of ITGBL1 Promotes Osteosarcoma Progression by Inhibiting Endoplasmic Reticulum Stress-Induced Autophagy. Advanced Science. 13(23), p.e15651.

Weber, F., Wagner, V., Rasmussen, S.B., Hartmann, R. and Paludan, S.R. 2006. Double-Stranded RNA Is Produced by Positive-Strand RNA Viruses and DNA Viruses but Not in Detectable Amounts by Negative-Strand RNA Viruses. Journal of Virology. 80(10), pp.5059–5064.

Winkler, L.L. and Kalejta, R.F. 2014. The 19S Proteasome Activator Promotes Human Cytomegalovirus Immediate Early Gene Expression through Proteolytic and Nonproteolytic Mechanisms K. Frueh, ed. Journal of Virology. 88(20), pp.11782–11790.

Wu, J., Wu, Y. and Lian, X. 2020. Targeted inhibition of GRP78 by HA15 promotes apoptosis of lung cancer cells accompanied by ER stress and autophagy. Biology Open., bio.053298.

Xu, L., Zhou, X., Peppelenbosch, M.P. and Pan, Q. 2015. Inhibition of hepatitis E virus replication by proteasome inhibitor is nonspecific. Archives of Virology. 160(2), pp.435–439.

Zhang, F., Xu, L.-D., Wu, S., Wu, Q., Wang, A., Liu, S., Zhang, Qian, Yu, X., Wang, B., Pan, Y., Huang, F., Neculai, D., Xia, B., Feng, X.-H., Shen, L., Zhang, Qi, Liang, T., Huang, Y.-W. and Xu, P. 2025. Proteasomal processing of the viral replicase ORF1 facilitates HEV-induced liver fibrosis. Proceedings of the National Academy of Sciences. 122(11), p.e2419946122.

Zhang, Y., Ma, X., Liu, C., Bie, Z., Liu, G., Liu, P. and Yang, Z. 2024. Identification of HSPD1 as a novel invasive biomarker associated with mitophagy in pituitary adenomas. Translational Oncology. 41, p.101886.

