## Supplementary Figures for "The proteasome and ERAD pathways can regulate hepatitis E virus pORF1 proteostasis and genome replication"

The authors have no potential conflicts of interest.

**Figure S1**

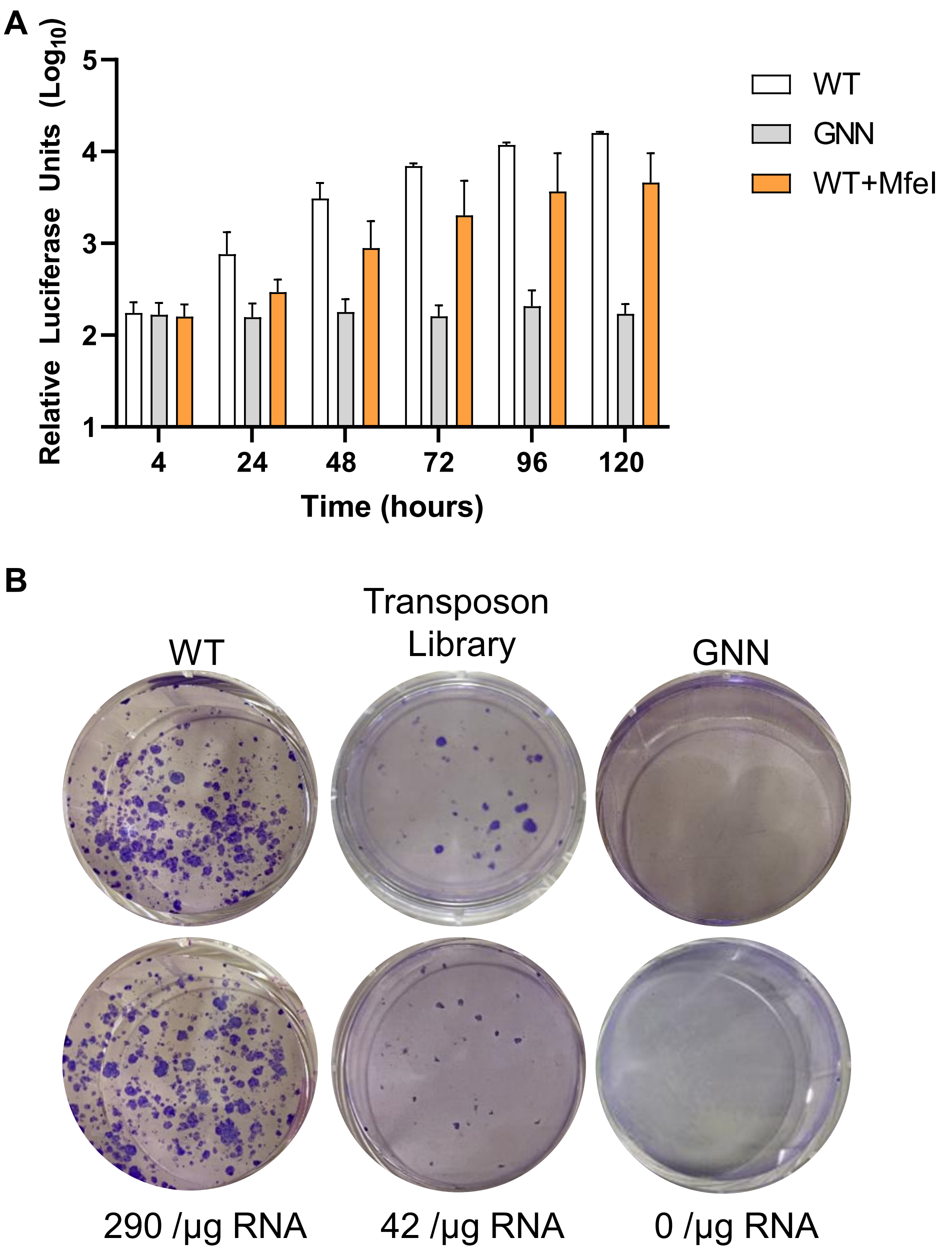


**Figure S1. Selection of transposon insertions in Huh7 cells.** (A) Huh7 cells were electroporated with SGR-G1-nLuc RNA containing an additional silent MfeI restriction site (WT+MfeI) together with wild-type (WT) and replication-defective (GNN) control SGR RNAs. Cells were harvested over 120 h post-electroporation and luciferase activity was determined. Data shown represent $\log_{10}$ of mean relative luciferase activity (n = 3 +/- SEM). (B) Huh7 cells were electroporated with SGR-G1-NPTII transposon library alongside wild-type (WT) and replication defective (GNN) control RNAs. Cells were selected with G418 for 14 days. Colony counts are shown as an average across two experiments and per µg of SGR RNA.

**Figure S2**

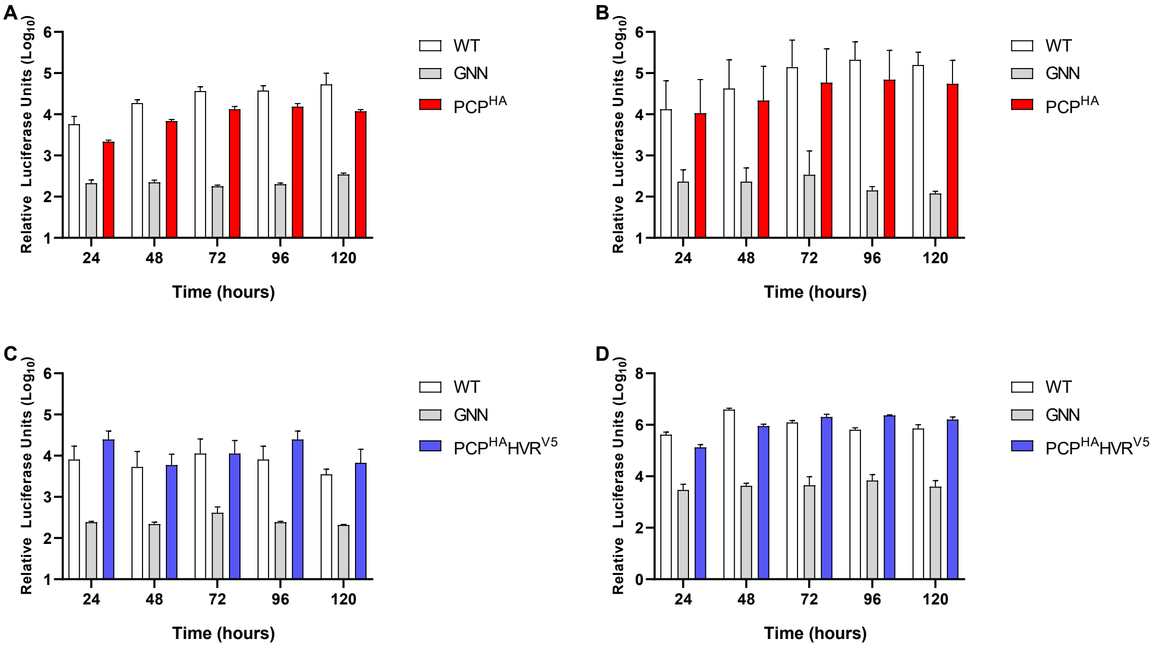


**Figure S2. Replication of epitope-tagged HEV SGRs**. Huh7 cells were electroporated with (A) SGR-G1-PCP$^{HA}$ or (B) SGR-G3-PCP$^{HA}$ alongside wild-type (WT) and replication-defective (GNN) controls. Cells were harvested at 24 h intervals across a 120 h timecourse and luciferase activity determined. Data shown as $\log_{10}$ of mean relative luciferase units (n = 3 ± SEM). Huh7 cells (C) and SK-RST cells (D) were electroporated with the dual-tagged SGR-G3-PCP$^{HA}$HVR$^{V5}$ alongside wild-type (WT) and replication-defective (GNN) controls. Cells were harvested at 24 h intervals across a 120 h time course, and luciferase activity was determined. Data are shown as $\log_{10}$ of mean relative luciferase units (n = 3 ± SEM).

**Figure S3**

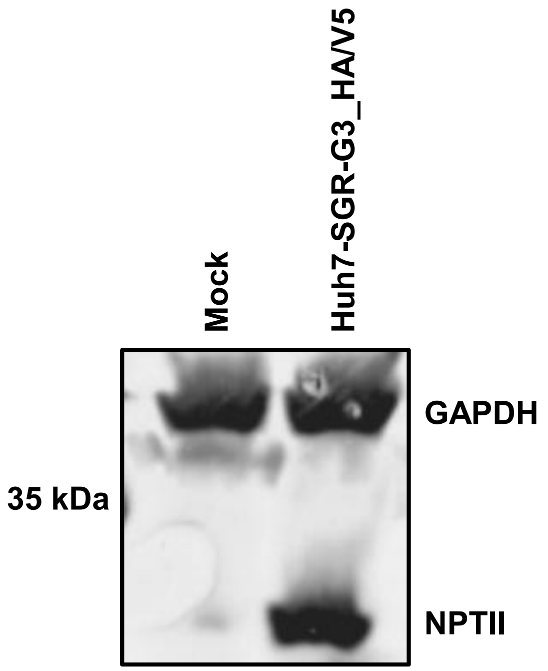


**Figure S3. Western blot analysis for expression of NPTII in the Huh7-SGR-G3-HA/V5 stable cell line.** Western blot analysis of lysates from naïve Huh7 and Huh7-SGR-G3-HA/V5 cells. Membranes were probed for anti-NPTII, and anti-GAPDH as a loading control.

**Figure S4**

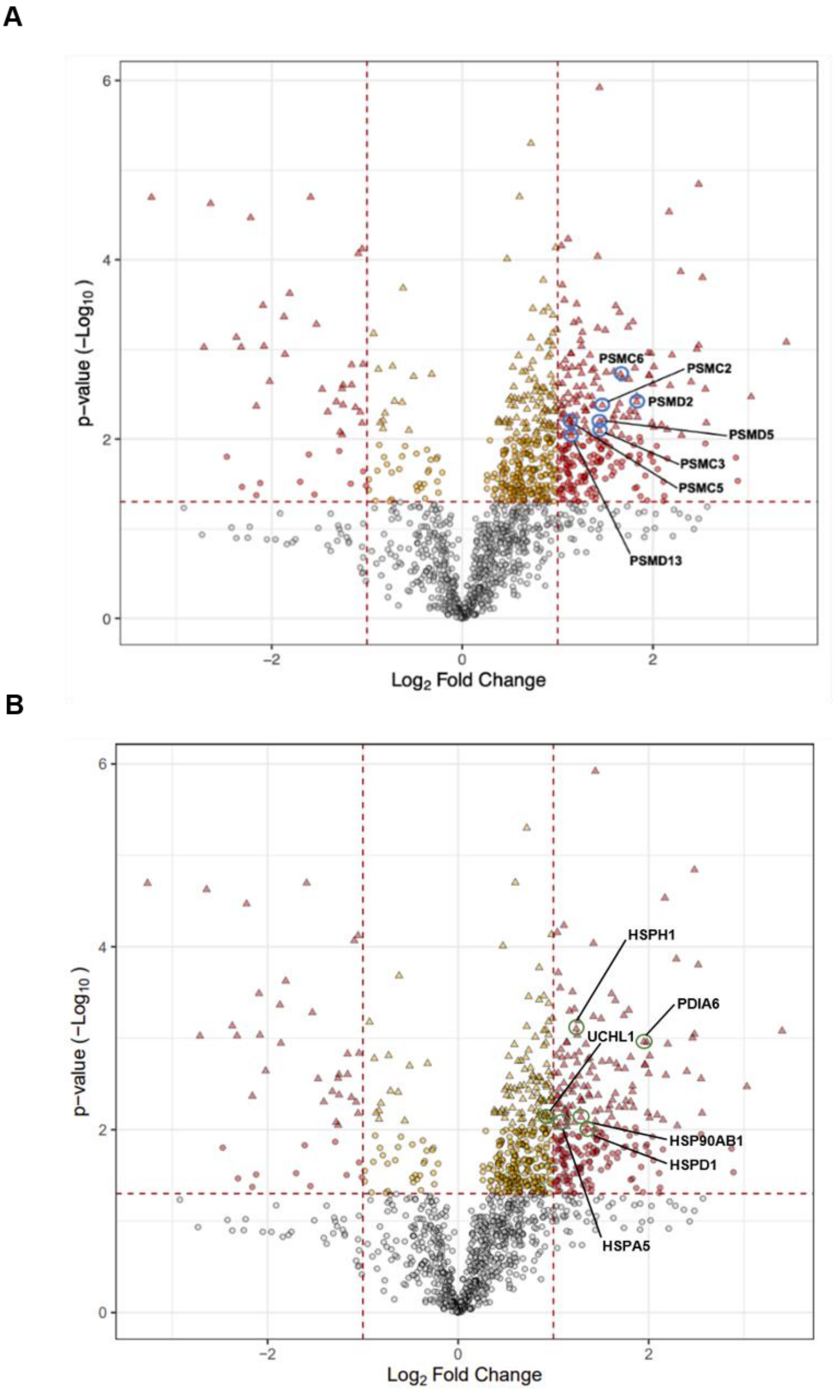


**Figure S4. Volcano plot showing differential protein enrichment identified by TMT-based LC-MS analysis**. Statistical analysis of proteins isolated within an anti-HA IP from Huh7-SGR-G3-HA/V5 stable cells compared with naïve Huh7 cells. Three biological replicates per condition were analysed. The x-axis represents $\log_2$ fold change, and the y-axis represents ($-\log_{10}$) adjusted p-value. Each point (triangle) corresponds to a protein identified in the anti-HA IP. Proteasome-related proteins (A, blue circles) and ERAD-related proteins (B, green circles) are highlighted.

**Figure S5**

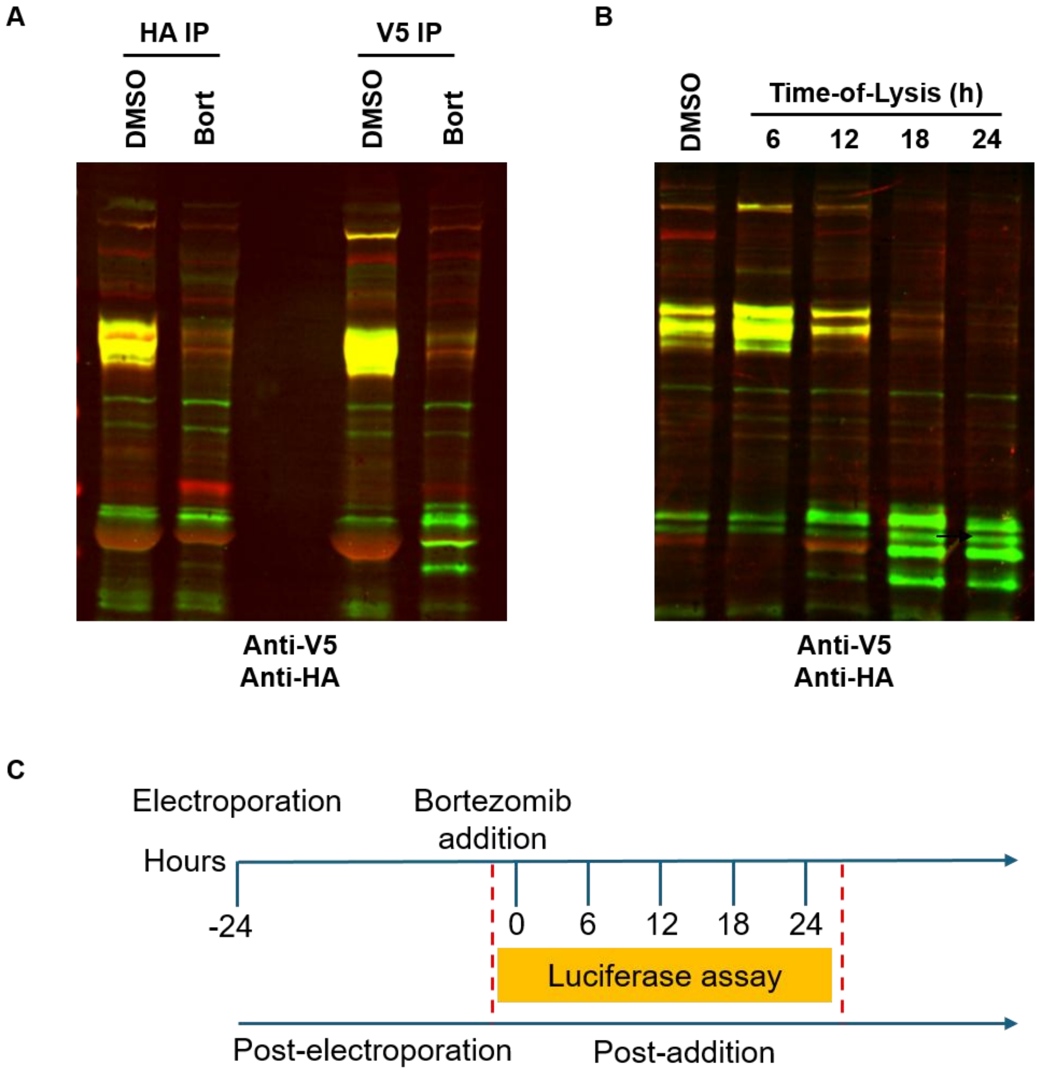


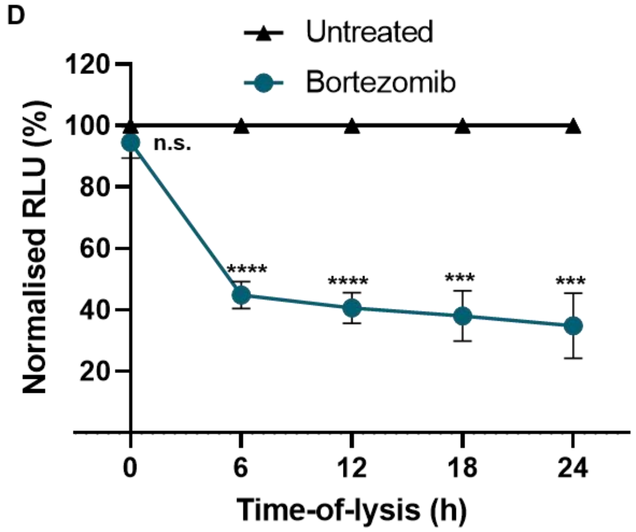


E

| | Untreated (RLU) | Bortezomib (RLU) |
|---|---|---|
| 0 | 17603 | 16517 |
| 6 | 22057 | 9830 |
| 12 | 30673 | 12635 |
| 18 | 40300 | 15412 |
| 24 | 40615 | 14397 |

**Figure S5. Bortezomib treatment reduces HEV replication and disrupts pORF1 proteolysis.** (A) Merged anti-V5 (green) and anti-HA (red) western blot analysis from bortezomib- and DMSO-treated Huh7-SGR-G3-HA/V5 cells (B) Merged anti-V5 and anti-HA western blot analysis from anti-V5 IP samples subject to timecourse analysis. (C) Schematic to illustrate experiment performed whereby Huh7-SGR-G3-HA/V5 electroporated cells were subject to DMSO or bortezomib treatment. (D) Time-course analysis of HEV ORF1 stability and replication under bortezomib treatment. Huh7 cells were electroporated with G3 HEV SGR and then treated with 1000 nM bortezomib or DMSO. Cell supernatants were collected at 6, 12, 18, and 24 h post-treatment, and luciferase activity was measured. Data shown represents a normalisation of mean relative luciferase activity to each individual untreated time point (n = 3 +/- SEM). (E) Tabulated mean RLU for three independent experimental repeats for supernatants harvested at 0, 6, 12, 18 and 24 hours post DMSO (black) or bortezomib (blue) addition.
